# Control of contextual memory through interneuronal α5-GABA_A_ receptors

**DOI:** 10.1101/2022.03.03.482912

**Authors:** Mengwen Zhu, Alifayaz Abdulzahir, Mark G Perkins, Chan C Chu, Bryan M Krause, Cameron Casey, Richard Lennertz, David Ruhl, Harald Hentschke, Rajasekar Nagarajan, Edwin R Chapman, Uwe Rudolph, Michael S Fanselow, Robert A Pearce

## Abstract

γ-aminobutyric acid type A receptors that incorporate α5 subunits (α5-GABA_A_Rs) are highly enriched in the hippocampus and are strongly implicated in control of learning and memory. Receptors located on pyramidal neuron dendrites have long been considered responsible, but here we report that their selective knockout from either interneurons (α5-i-KO) or pyramidal neurons (α5-pyr-KO) interferes with the ability of the general anesthetic etomidate to suppress contextual conditioning. Using Ca^2+^ imaging of CA1 pyramidal neuron activity in freely exploring mice to assess hippocampal function directly, we found that etomidate blocked the development of place cells and spatial engrams in wild type (WT) and α5-pyr-KO mice, but not in α5-i-KO mice. In addition, α5-i-KO mice developed weaker spatial engrams than WT mice under control conditions. These findings show that interneuronal α5-GABA_A_Rs serve a physiological role in promoting spatial learning, and that they mediate the suppression of hippocampus-dependent memory by etomidate.

## Introduction

Inhibitory synaptic transmission in the brain is mediated primarily by γ-aminobutyric acid type A receptors (GABA_A_Rs). This family of chloride-permeable ionotropic receptors is the target of a wide variety of clinically important agents, including sedative-hypnotics, anxiolytics, anticonvulsants, and general anesthetics ^1, 2^. Many of these agents also impair memory. Usually this is an undesired side-effect, but for general anesthetics it is a primary goal. Indeed, awareness with recall during anesthesia is recognized as a persistent problem ^3, 4, 5^, and it has been associated with dysphoria and post-traumatic stress disorder ^4, 6^. In studies of the mechanism of anesthetic-induced amnesia, etomidate has proved exceptionally useful, in part because it is highly selective for GABA_A_ receptors, but also because its physicochemical properties make it amenable to use both *in vivo* and *in vitro* ^7, 8^.

Etomidate potently blocks memory formation even at subanesthetic doses by targeting GABA_A_Rs that incorporate α5 subunits (α5-GABA_A_Rs) ^9, 10^. These receptors are highly concentrated in the hippocampus ^11, 12^, a brain structure that is essential for the formation of episodic memories ^13^. The well-established role of α5-GABA_A_Rs in learning and memory has prompted the development of agents that target α5-GABA_A_Rs to enhance cognition ^14, 15, 16, 17, 18^, with application to diverse disease states including Alzheimer’s disease, Down syndrome, autism, depression, and schizophrenia ^16, 19, 20, 21, 22, 23, 24, 25, 26^.

The generally accepted mechanism by which α5-GABA_A_Rs modulate memory involves receptors at extrasynaptic sites on pyramidal neurons ^27, 28, 29^. These receptors are located primarily at extrasynaptic sites on pyramidal neuron dendrites ^30^, and acting through either tonic ^31^ or slow phasic inhibition ^32^, α5-GABA_A_Rs are well suited to prevent the depolarization and NMDAR-mediated calcium entry that initiates synaptic plasticity. Accordingly, GABAergic drugs such as etomidate that are normally able to suppress long-term potentiation (LTP) and learning are ineffective in mice with a global knockout of α5 subunits (α5-gl-KO), or in mice administered α5-selective negative modulators ^9, 10^. However, recent studies indicate that α5-GABA_A_Rs are not restricted to pyramidal neurons ^33, 34^, raising the possibility that interneuronal α5-GABA_A_Rs might also play a role. Indeed, we found that selective knockout of α5 subunits from pyramidal neurons did not recapitulate the resistance to LTP suppression seen in α5-gl-KO mice ^35^, and that suppression of LTP and contextual fear conditioning were independent of GABA_A_Rs that incorporate β3 subunits, which underlie tonic and long-lasting phasic inhibition in pyramidal neurons ^36, 37^. These findings raised the possibility that α5-GABA_A_Rs on pyramidal neurons might not be the only targets mediating etomidate-induced amnesia. In the present study we therefore tested the contributions of α5-GABA_A_Rs on interneurons *versus* pyramidal neurons by studying the effects of etomidate on contextual fear conditioning as well as place cells and spatial engrams as neural correlates of spatial memory, in interneuron-selective α5-GABA_A_R knockout (α5-i-KO), pyramidal neuron-selective α5-GABA_A_R knockout (α5-pyr-KO) mice, and pseudo-WT (p-WT) mice.

## Results

### Cell type-specific elimination of α5-GABA_A_Rs

To generate mice lacking α5-GABA_A_Rs specifically in interneurons or pyramidal neurons, we crossed mice carrying a floxed α5 allele (Gabra5^tm2.1Uru^) with mice that express Cre recombinase under the control of the GAD2 promoter (GAD2^tm2(Cre)Zjh^) or CaMKIIa promoter (Tg(Camk2a-Cre)T29-1Stl) respectively (Fig. 1a). We showed previously by immunohistochemistry that α5-GABA_A_Rs in hippocampus are strongly reduced in α5-pyr-KO mice, to a level comparable to that of global α5-GABA_A_R knockout mice (α5-gl-KO), consistent with a predominance of α5-GABA_A_Rs in pyramidal neurons ^35^. Western blot analysis of α5-i-KO mice similarly showed that p-WT mice and α5-i-KO mice expressed α5 subunits at levels indistinguishable from C57BL/6J (WT) mice (Fig. 1b), again consistent with the predominant expression of α5-GABA_A_Rs in pyramidal neurons. To test whether the GAD2-Cre promoter might lead to loss of α5-GABA_A_Rs in astrocytes, which also express GABA_A_Rs and synthesize and release GABA ^38^, we tested for co-expression of the GAD2-Cre-driven tdTomato reporter and GFAP-driven GFP (Fig. 1c). We found complete segregation of the markers (Fig. 1d), further supporting the restriction of α5 subunit knockout to interneurons but not glia in α5-i-KO mice. Finally, to directly demonstrate a reduction in α5-GABA_A_R protein in interneurons, we examined co-localization of α5 subunits and GAD2 in WT, α5-i-KO, α5-pyr-KO, and α5-gl-KO mice. We found that α5-GABA_A_Rs and GAD2 were co-localized on many interneurons in *stratum oriens* and *stratum radiatum* in WT and α5-pyr-KO mice, but co-localized markers were entirely absent in α5-i-KO and α5-gl-KO mice (Fig. 1e). Together, these findings demonstrate that α5-GABA_A_Rs were successfully eliminated in a selective fashion in α5-i-KO and α5-pyr-KO mice.

**Fig. 1.**
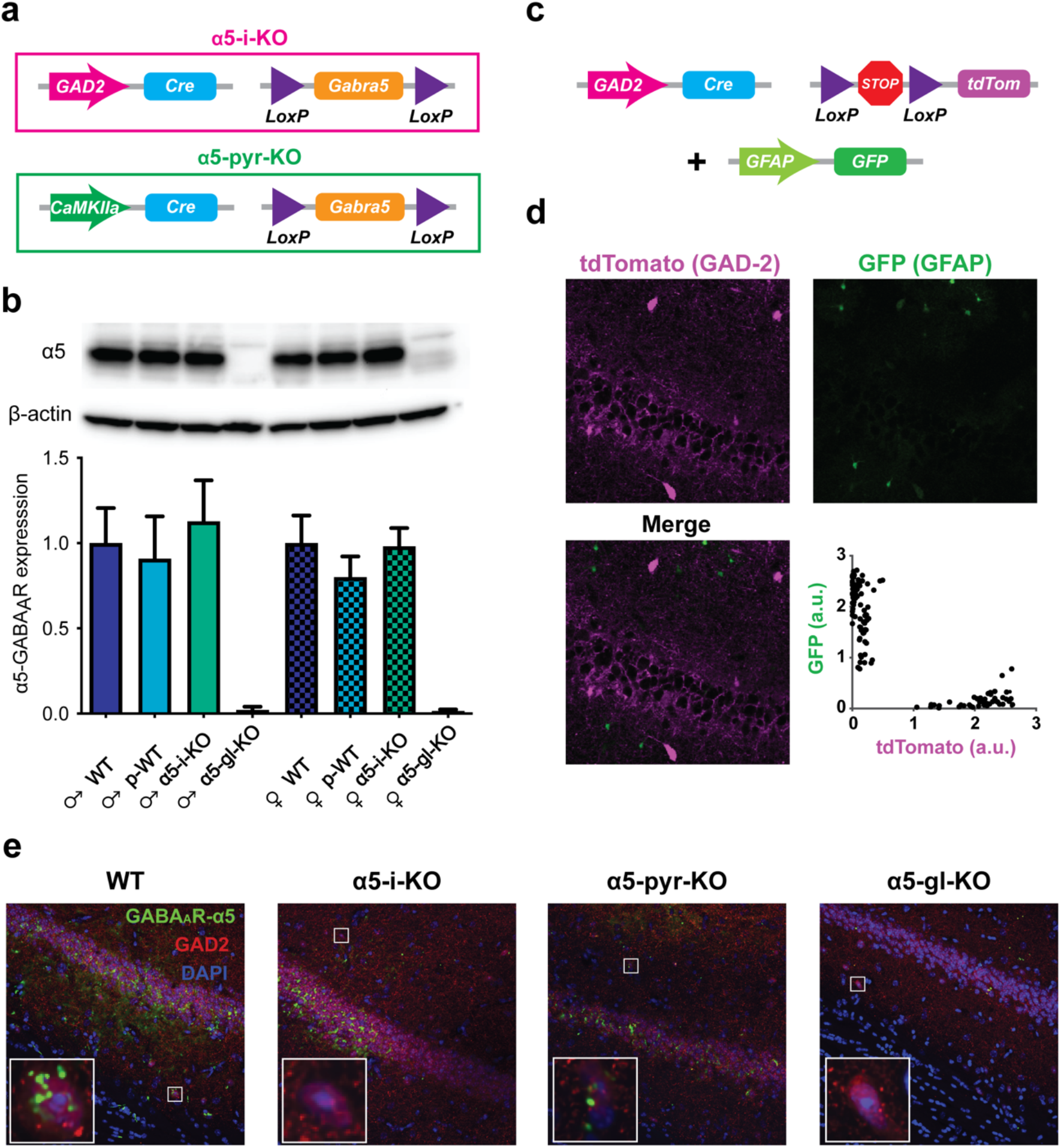
Specific elimination of α5-GABA_A_Rs from interneurons vs. pyramidal neurons. **a** Genetic strategies used to eliminate α5-GABA_A_Rs from interneurons (top) or pyramidal neurons (bottom). For α5-i-KO mice, Cre recombinase was driven by the interneuron-specific GAD2 promoter to target the floxed Gabra5 gene. For α5-pyr-KO mice, Cre was driven by the pyramidal neuron-specific CaMKIIa promoter. **b** Western blot analysis of α5-GABA_A_R (55 kDa) expression in male and female WT, p-WT, α5-i-KO, and α5-gl-KO mice. Expression levels were normalized to β-actin (42 kDa) and quantified by densitometry. Note that receptor levels were unchanged in p-WT and α5-i-KO mice versus WT mice, consistent with maintained expression in pyramidal neurons. **c** Genetic strategy used to test the overlap between Cre recombinase expression in interneurons (Cre-dependent td-Tomato expression) versus astrocytes (GFP driven by the astrocyte-specific GFAP promoter). **d** Confocal imaging of transgene expression. Fluorescence levels (in a.u.—arbitrary units) of GAD2-Cre driven td-Tom expression and GFAP-driven GFP expression were highly segregated, demonstrating a lack of GAD2-driven Cre recombinase in astrocytes. **e** Immunohistochemical verification of loss of α5-GABA_A_Rs from interneurons vs. pyramidal neurons. Insets show representative interneurons, with loss of α5-GABA_A_R staining from α5-i-KO and α5-gl-KO mice.

### α5-i-KO and α5-pyr-KO mice resist the suppression of contextual conditioning by etomidate

To test the role of α5-GABA_A_Rs expressed on interneurons and pyramidal neurons in etomidate’s suppression of hippocampus-dependent memory, we employed the Context Preexposure Facilitation Effect (CPFE) learning paradigm (Fig. 2a) – a variant of contextual fear conditioning that separates the contextual learning and aversive phases of conditioning ^39, 40, 41, 42^. This paradigm takes advantage of the “immediate shock deficit” ^43^, wherein mice that are shocked immediately after they are placed in a novel context fail to associate the context and shock, whereas mice that were pre-exposed to the context on the preceding day recall the context rapidly and successfully associate it with the shock ^44, 45^. Because the drug is present only during contextual learning (Day 1) and not when the shock is administered (Day 2), any observed change in freezing behavior (Day 3) can be attributed to a failure to learn the context rather than an attenuation of the shock’s effectiveness.

**Fig. 2.**
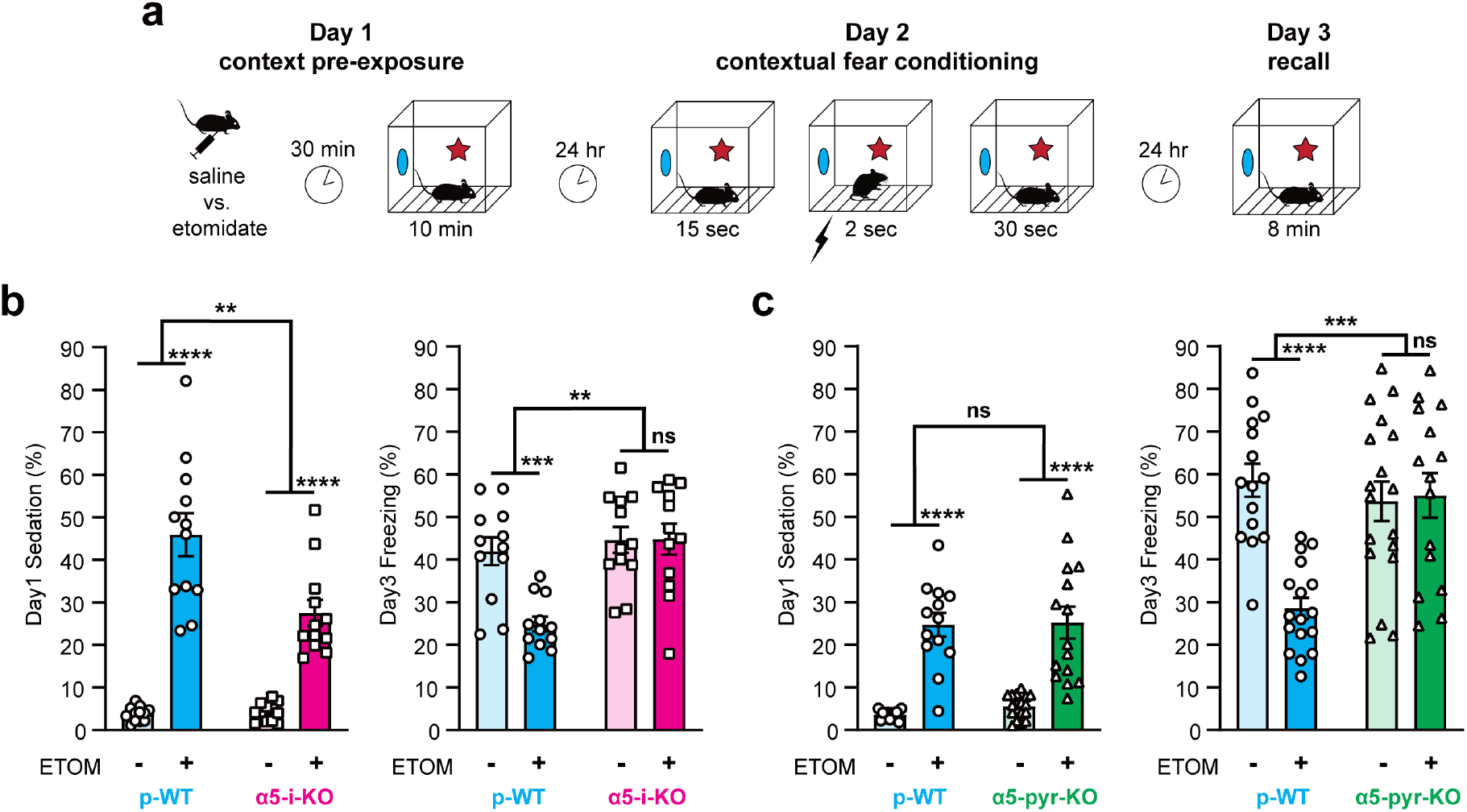
α5-i-KO and α5-pyr-KO mice resist the suppression of contextual learning by etomidate. **a** Context Preexposure Facilitation Effect (CPFE) paradigm. **b** Effect of etomidate in α5-i-KO and p-WT mice. Etomidate (7 mg/kg IP) sedated both gentoypes, p-WT more than α5-i-KO mice (left), but impaired memory in only p-WT mice (right). *p < 0.05, **p < 0.01, ***p < 0.001, ****p < 0.0001. **c** Effect of etomidate in α5-pyr-KO and p-WT mice. Etomidate sedated both gentoypes equally, but impaired memory in only p-WT mice (right). *p < 0.05, **p < 0.01, ***p < 0.001, ****p < 0.0001.

There were no baseline differences between α5-i-KO mice and their p-WT littermates (p-WT_i_) in anxiety, as indicated by open arm entries or open arm time in the elevated plus maze test, or in pain sensitivity, as indicated by time to withdrawal in the hot plate test (Supplementary Fig. 1). Etomidate (7 mg/kg IP, 30 min before Day 1) did produce a greater level of sedation (Day 1) in p-WT_i_ compared to α5-i-KO mice (Fig. 2b, *left*; drug x genotype, F(1,44) = 10.04, p = 0.0028), but on average both genotypes spent more than one-half of their time actively exploring the novel context during the context preexposure. Most importantly, etomidate differentially suppressed freezing (Day 3) in α5-i-KO versus p-WT_i_ mice (Fig. 2b, *right*; drug x genotype, F(1,44) = 8.09, p = 0.0067), with a significant suppression of freezing in p-WT_i_ mice (t(44) = 3.96, p < 0.001) but not in α5-i-KO mice (t(44) = 0.0365, p = 0.99). These results demonstrate that α5-GABA_A_Rs on interneurons are essential for suppression of contextual fear conditioning by a sedative dose of etomidate.

To test whether α5-GABA_A_Rs on pyramidal neurons also contribute to etomidate’s suppression of contextual fear conditioning, we similarly performed CPFE experiments in α5-pyr-KO mice and their p-WT littermates (p-WT_pyr_). Etomidate produced equal levels of sedation (Day 1) in p-WT_pyr_ and α5-pyr-KO mice (Fig. 2c, *left*; drug x genotype, F(1,61) = 0.19, p = 0.66). Again, etomidate differentially suppressed freezing (Day 3) in the two genotypes (Fig. 2c, *right*; drug x genotype, F(1,61) = 14.17, p<0.001), with significant suppression in p-WT_pyr_ mice (t(61) = 5.06, p < 0.0001) but not in α5-pyr-KO mice (t(61)=0.233, p = 0.97). This finding indicates that etomidate also engages pyramidal neuron α5-GABA_A_Rs to block memory, consistent with previous experiments linking these receptors on CA1 pyramidal neurons specifically to the control of spatial memory ^46^.

### Etomidate suppresses place cell formation via interneuronal α5-GABA_A_Rs

The hippocampus supports contextual memory by forming a cognitive map of the environment ^47, 48^. Spatially modulated firing of individual cells, referred to as ‘place cells’, contribute to that map ^49, 50^. Their spatial specificity can be stable for periods of days or even weeks in a given environment ^51, 52^, but they undergo ‘remapping’ when the animal is placed in a new environment ^53, 54^. Although not every place cell is seen again upon re-exposure to the same environment, when it is reactivated, its place field is retained ^55^. Therefore, as an ensemble, the formation and re-activation of place cells can serve as a ‘readout’ of the encoded contextual memory, stable over a period of weeks ^56^.

To test the influence of etomidate on this process, and whether it depends on α5-GABA_A_Rs on interneurons and/or pyramidal neurons, we used *in vivo* Ca^2+^ imaging in freely moving mice to assess the spatially modulated firing characteristics of dorsal CA1 pyramidal neurons in p-WT, α5-i-KO, and α5-pyr-KO mice as they explored novel contexts two to three times per week for up to ten weeks ^57^. To capture Ca^2+^ signals, we injected virus carrying the genetically encoded calcium indicator GCaMP6f driven by the CaMKIIa promoter (AAV1-CaMKIIa-GCaMP6f) into the dorsal hippocampus of 16 mice (7 p-WT, 5 α5-i-KO, 4 α5-pyr-KO). Two to three weeks later, a gradient refractive index (GRIN) lens and baseplate were attached to the skull, to which was affixed a miniature endoscope (nVoke2, Inscopix, Palo Alto, CA). Beginning ~4-6 weeks after virus injection, transient fluorescent signals were observed in up to ~800 individual CA1 pyramidal neurons (Fig. 3a and Supplementary Fig. 2). Over the course of a 10-minute period of free exploration in a novel environment, overall Ca^2+^ event rates averaged ~0.1 sec^−1^, similar to previous reports ^55, 58^.

**Fig. 3.**
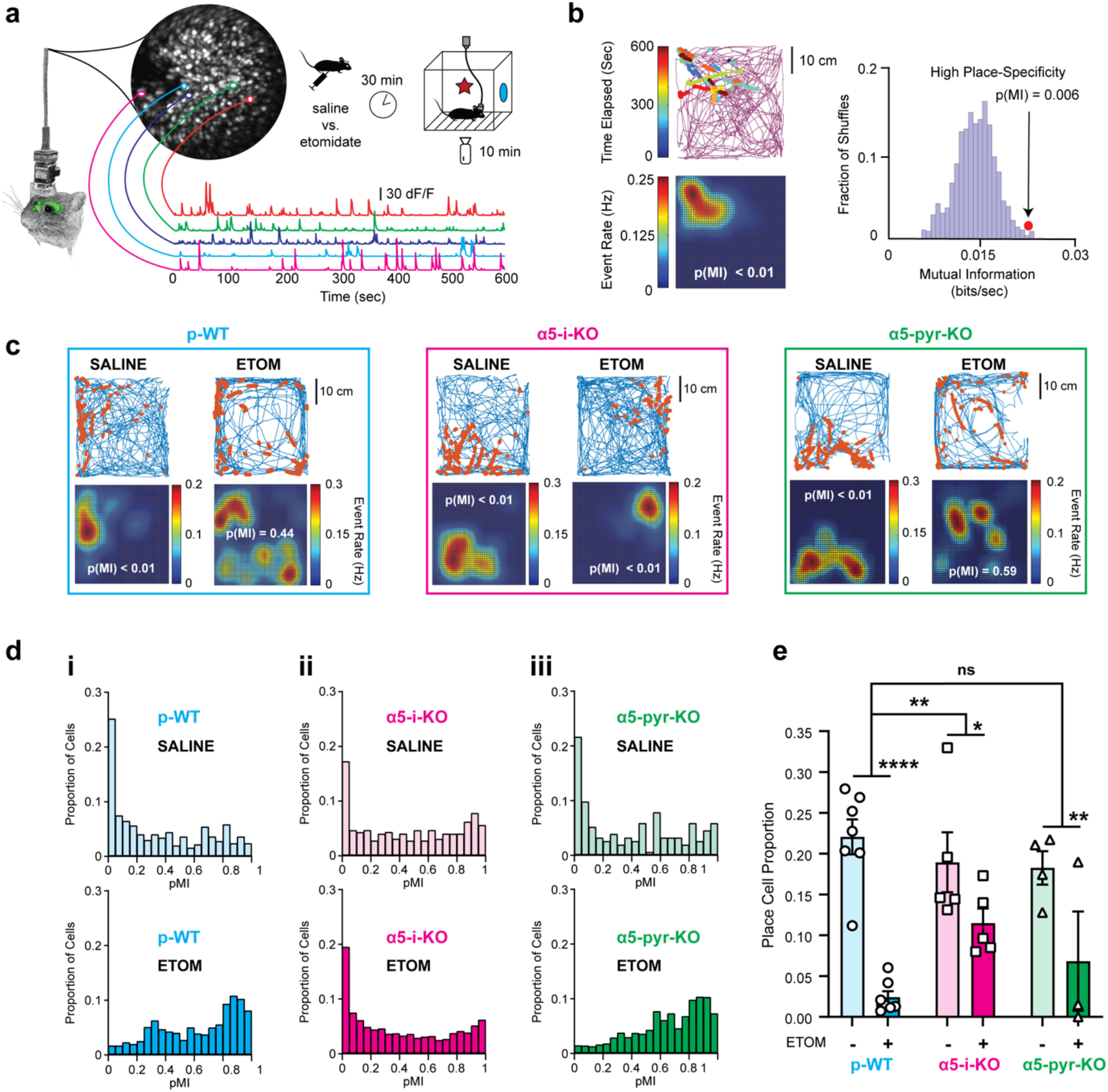
Suppression of place cell formation by etomidate is attenuated in α5-i-KO but not α5-pyr-KO mice. **a** Hippocampal dorsal CA1 pyramidal neuron activity was captured using a head-mounted miniaturized endoscope to monitor AAV1-CaMKIIa-GCaMP6f fluorescence. Contours of five cells are overlayed on the maximum projected image from a 10-minute recording; their associated fluorescence traces are shown below. Upper right: experimental paradigm used to evaluate place cell formation. Ca^2+^ dynamics and arena location were captured simultaneously. **b** Establishing the place-specific firing of a cell. Upper left: event dot map shows the location of the mouse when Ca^2+^ events were detected. The continuous purple line shows the track of the mouse in the experimental arena during the entire 10 minutes. Each string of dots represents a single Ca^2+^ event distributed along the rising phase of the Ca^2+^ transient, and the number of dots in each string reflects the event’s adjusted amplitude. The string color designates the time at which the event occurred. Lower left: gaussian-smoothed event rate map of the corresponding event dot map above. Warmer colors represent higher event rates. Right: mutual information (MI) between mouse position and firing rate for the cell shown on the left *versus* its time-shuffled null distribution. In this example, the probability of the observed MI (red dot) falling within the null distribution was p(MI)=0.006, which classifies the cell as a place cell. **c** Examples of event dot maps and their derived event rate maps for representative cells after mice had been administered saline or etomidate (7 mg/kg). Blue lines show the track of the mouse and orange dots represent calcium events. Left: place-specific firing of a cell in a p-WT mouse under control conditions (p(MI) < 0.01) but not after etomidate (p(MI) = 0.44). Middle: place-specific firing of a cell from an α5-i-KO mouse after saline (p(MI) < 0.01) and also after etomidate (p(MI) < 0.01). Right: similar to p-WT, place-specific firing in a α5-pyr-KO mouse after saline (p(MI < 0.01) but not after etomidate (p(MI) = 0.59). **d** p(MI) distributions of all cells from six representative sessions after saline or etomidate (7 mg/kg). Left (i): the left-skewed distribution of p(MI) (e.g. more place-specific cells) of a p-WT mouse under saline shifts to the right (e.g. cells losing place specificity) under etomidate. Middle (ii): the distribution of p(MI) retains its left-skewed shape despite of the presence of etomidate in an α5-i-KO mouse. Right (iii): similar to p-WT, the p(MI) distribution of an α5-pyr-KO mouse shifts to the right under etomidate. **e** The effect of etomidate 7mg/kg on place cell formation summarized for all three genotypes. Etomidate suppression of place cell formation was attenuated in α5-i-KO mice but not α5-pyr-KO mice. There was no influence of genotype on place cell proportions after saline (F(2, 12.45) = 0.31, p = 0.74) (*p < 0.05, **p < 0.01, ****p < 0.0001).

To quantify place-specific firing of individual neurons, we simultaneously monitored the position of the mouse using video tracking software (Noldus EthoVision XT15) together with endoscopic fluorescent report of cellular activity (Inscopix Data Acquisition Software 2019). Fig. 3b shows an example of one such experiment. The track of the mouse over the 10-min exploration is shown by the purple line, with dots superimposed over the rising phase of detected calcium events (Fig. 3b, *upper left*). This information was used to create a map of event rate as a function of position (Fig. 3b, *lower left*). To quantify the place-specificity of Ca^2+^ dynamics of a specific cell, we calculated the mutual information (MI) between the cell’s event map and mouse’s spatial location map and compared it to a null distribution of MI values obtained from 1000 time-shuffled event maps (Fig. 3b, *right* and Supplementary Fig. 3A). Using the criterion that the probability of the observed MI value must fall within the top 5% of the null distribution (i.e. p(MI) < 0.05) for a cell to be considered a “place cell”, we found that in p-WT mice under control conditions, 22 ± 2% of all active cells (≥5 Ca^2+^ events over a 10-minute recording) qualified as place cells, consistent with previous reports using this recording method ^55^.

To determine whether an amnestic dose of etomidate altered place cell formation during a 10-minute exploratory session in a novel environment – matching the Day 1 (context preexposure) of CPFE experiments (Fig. 2a) – we compared the number of place cells seen in mice that had received an IP injection of saline or 7 mg/kg etomidate 30 minutes prior to placing the mouse in the experimental arena. Examples of activity tracks and Ca^2+^ event rate maps, with associated p(MI) values for individual neurons, are shown in Fig. 3c, for p-WT, α5-i-KO, and α5-pyr-KO mice when saline or 7 mg/kg etomidate were administered. As it did in CPFE experiments, etomidate produced a moderate level of sedation in all genotypes, as indicated by the fraction of time the mouse actively explored (≥1 cm/sec) the experimental arena, but its effect was significantly stronger in p-WT and α5-pyr-KO than in α5-i-KO mice (Supplementary Fig. 4). Etomidate also produced a slight but significant dose-dependent reduction in event rate in p-WT and α5-pyr-KO mice, but not in α5-i-KO mice (Supplementary Fig. 5). Most strikingly, in p-WT mice, etomidate shifted the distribution of p(MI) values from one heavily skewed toward low p(MI) values under control conditions (Fig. 3d, i, top) toward the right (Fig. 3d, i, bottom), essentially eliminating the proportion of cells with p(MI)<0.05 (i.e. place cells). By contrast, in α5-i-KO mice, the distribution and place cell proportion were not different than p-WT under control conditions, but the effect of etomidate was greatly attenuated, and the proportion of place cells was largely maintained (Fig. 3d, ii). Etomidate’s effect on α5-pyr-KO mice was not different than on p-WT mice (Fig. 3d, iii).

A summary of the place cell proportions seen in all mice administered saline or 7 mg/kg etomidate is presented in Fig. 3e. On average, etomidate reduced the fraction of place cells from 22 ± 2% (saline) to 2 ± 1% in p-WT mice (t(46.23) = −8.87, p < 0.0001), whereas in α5-i-KO mice the place cell fraction was reduced from 19 ± 5% (saline) to 12 ± 2% (t(25.99) = −2.56, p = 0.016), a significantly weaker effect (drug x genotype, t(103.69) = 2.74, p = 0.0071). In α5-pyr-KO mice, etomidate reduced the place cell proportion from 18 ± 2% (saline) to 7±5% (t(21.17) = −3.11, p = 0.0028), an effect that was not significantly different than p-WT (drug x genotype, t(114.23) = 1.065, p = 0.29).

To establish the dose dependence of place cell suppression, we carried out this same type of experiment using doses of 2, 4, 6, and 8 mg/kg etomidate. The findings mirrored those presented above, with strong dose-dependent suppression of place cell proportion in p-WT and α5-pyr-KO mice and resistance to this effect in α5-i-KO mice (Supplementary Fig. 6). The finding that place cell suppression depends on α5-GABA_A_Rs on interneurons mirrors our CPFE results (Fig. 2), supporting a causal link between etomidate’s interference with place cell formation and hippocampus-dependent contextual learning. By contrast, the sensitivity of α5-pyr-KO mice to etomidate’s suppression of place cells suggests that freezing in the CPFE experiments may have reflected a learning strategy that does not rely in the same way on spatially modulated firing.

### Etomidate suppresses spatial engrams via interneuronal α5-GABA_A_Rs

Once formed, place cells can remain stable for days to weeks, displaying the same position-specific activity upon repeated exposure to the same environment ^56^, thereby forming a memory trace, or ‘spatial engram’, of the environment. Cells that do not meet the relatively strict criterion of p(MI)<0.05 also carry some spatial information and contribute to the spatial engram ^56, 59, 60^. Therefore, to test the influence of etomidate on spatial engram stability, we measured the similarity between firing patterns during a mouse’s initial exposure to a novel environment in ‘Session 1’ (S1) versus its re-exposure to the same environment 4 or 24 hours later in ‘Session 2’ (S2), using event rate maps from all cells that were active during both sessions (Fig. 4a). We quantified the similarity of spatially modulated calcium dynamics in two ways: first, on a cell-by-cell basis, by computing the Pearson’s correlation coefficient (PCC) between rate maps (RMs) formed by same cells in S1 and S2 (Supplementary Fig. 3B); and secondly, on an ensemble basis, by computing the PCC between population vectors (PVs) of firing rates of all cells at each position in the arena (divided into 15-by-15 pixels) (Supplementary Fig. 3C). Results for these two methods were essentially identical, so here we present RM_corr_ results; PV_corr_ results are presented in supplementary figures (Supplementary Fig. 7).

**Fig. 4.**
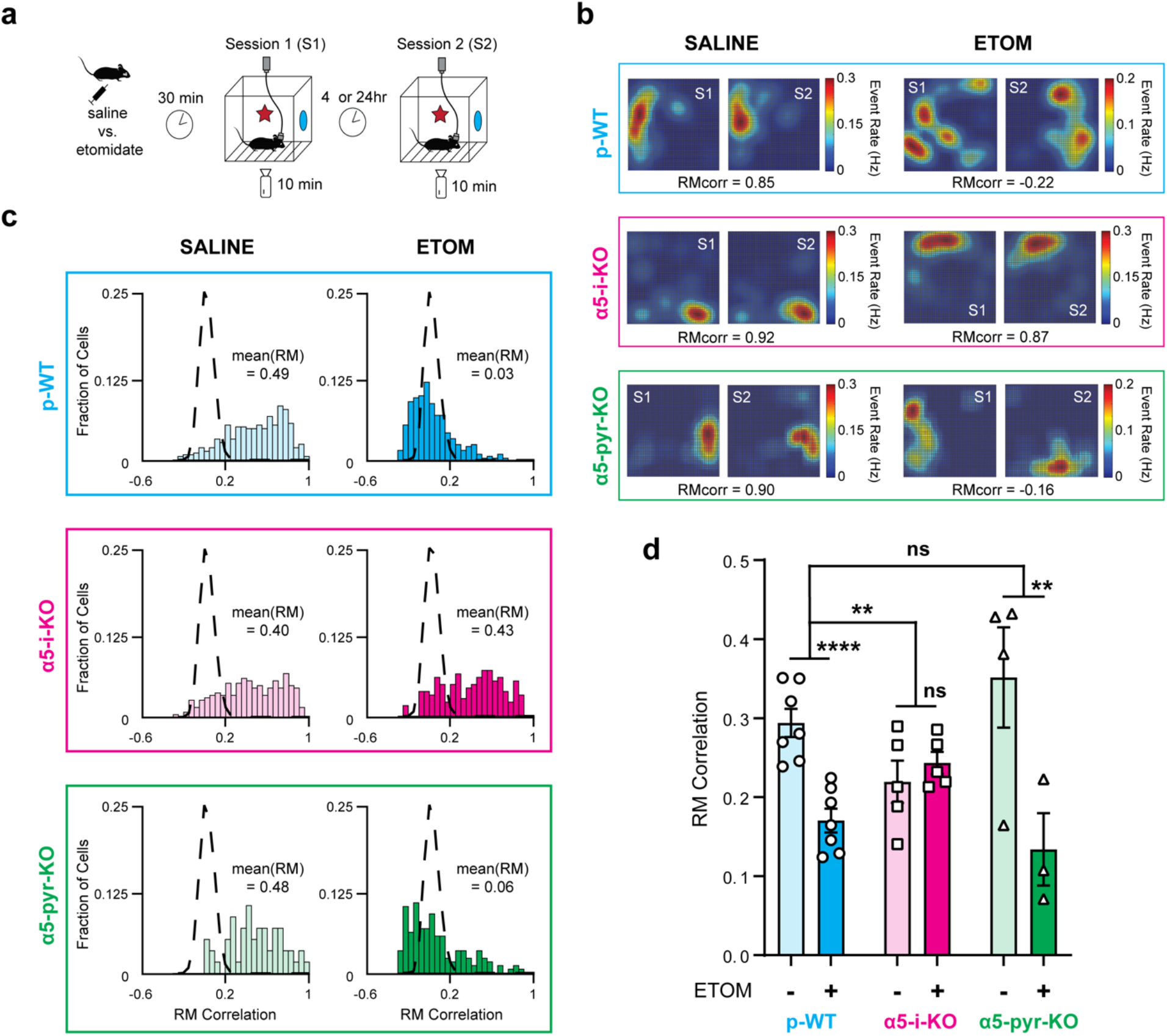
Etomidate suppresses hippocampal spatial engram stability in p-WT and α5-pyr-KO mice but not in α5-pyr-KO mice. **a** Experimental paradigm used to assess spatial engram formation. The same mice used to assess place cell formation (Fig. 3) were re-exposed to the same contexts in a second session. Saline or etomidate was administered only before the first session. **b** Representative examples of event rate maps for three individual neurons from p-WT (top), α5-i-KO (middle), and α5-pyr-KO (bottom) mice that were administered saline (left) or 7mg/kg etomidate (right). Their associated rate map (RM) correlation between S1 and S2 are shown below. Under control conditions (saline) cells retained their place specificity. After etomidate, cells in p-WT and α5-pyr-KO, but not α5-i-KO mice, underwent remapping. **c** Distributions of RM correlations from six paired recording sessions in p-WT (top), α5-i-KO (middle), and α5-pyr-KO (bottom) mice that were administered saline (left) or 7mg/kg etomidate (right). Left: under control conditions, distributions fell substantially to the right of the location-shuffled RM null distributions (dashed lines) in all three genotypes, revealing the presence of stable spatial engrams. Right: etomidate caused the distributions to be shifted toward the null distributions in p-WT and α5-pyr-KO mice but not in α5-i-KO mice. **d** Summarized mean RM correlation for all three genotypes under saline and etomidate 7mg/kg conditions. Etomidate strong impaired spatial engram stability in both p-WT and α5-pyr-KO mice but not in α5-i-KO mice. (**p < 0.01, ****p < 0.0001)

Representative examples of event rate maps and their associated RM correlation values (RM_corr_) from paired sessions are shown in Fig. 4b. These cells exemplified the activity patterns we observed: under control conditions (saline) for all three genotypes, cells tended to fire in the same spatial locations when mice were returned to the arena (Fig. 4b, *left*). In p-WT and α5-pyr-KO mice given 7mg/kg etomidate 30 minutes prior to S1, cells globally ‘remapped’ to new locations, but α5-i-KO mice maintained consistent spatial activity patterns (Fig. 4b, *right*). This resistance of α5-i-KO mice to etomidate was evident in the distributions of RM_corr_ values of all cells seen in both sessions (Fig. 4c): RM_corr_ values fell largely outside the location-shuffled null distribution (dashed lines) when mice were administered saline, indicating that they had formed stable spatial maps under control conditions (Fig. 4c, *left*); in contrast, 7mg/kg etomidate shifted RM_corr_ distributions toward null distribution in p-WT and α5-pyr-KO mice, but not in α5-i-KO mice (Fig. 4c, *right*).

Combined results from all such paired recording experiments in all genotypes, with saline or 7mg/kg etomidate administered prior to S1, are presented in Fig. 4d. Etomidate significantly reduced RM_corr_ in p-WT (t(45.69) = −5.20, p < 0.0001) and α5-pyr-KO mice (t(26.04) = −2.98, p = 0.0062), but not in α5-i-KO mice (t(37.31) = 0.727, p = 0.472). This drug effect was significantly different in α5-i-KO mice versus p-WT (drug x genotype; t(113.20) = 3.095, p = 0.0025) but not in α5-pyr-KO mice (t (108.08) = −0.628, p = 0.53). Interestingly, the average RM_corr_ for α5-i-KO mice (0.22 ± 0.03) under saline control condition was significantly lower than that for p-WT mice (0.29 ± 0.02; t(59.19) = −2.26, p = 0.027), indicating that α5-GABA_A_Rs on interneurons play a physiological role in promoting spatial memory formation and/or stability. Similar experiments using etomidate doses of 2, 4, 6, and 8 mg/kg revealed dose-dependent reductions in RM_corr_ and PV_corr_ that were attenuated in α5-i-KO but not in α5-pyr-KO mice (Supplementary Figs. 7, 8). These results show that in addition to interfering with place cell formation, etomidate suppresses spatial engram formation and/or stability through α5-GABA_A_Rs on interneurons but not pyramidal neurons.

The analysis described above relies on similarities between spatially modulated firing patterns to establish the presence of a ‘spatial engram’. However, the traditionally recognized engram is based not on spatially modulated firing but on overall firing rates during the entire period of exploration (learning), high enough to activate early immediate genes (IEGs – cFos, Egr1, ARC, PNAS4) ^61, 62, 63^. To test whether a similar phenomenon might be present in the Ca^2+^ events that we measured here, we examined the effect of etomidate on the probability that a cell seen in S1 would also be active in S2 (cell recurring probability). We found no significant effects of etomidate in any of the three genotypes, though there was a trend toward lower cell recurring probability in p-WT and α5-pyr-KO mice but not in α5-i-KO mice (Supplementary Fig. 9). We conclude that IEG-defined engrams are distinct from spatial engrams based on overall Ca^2+^ event rates.

### Hippocampal spatial engrams are sensitive to local contextual cues and their representations of distal cues slowly evolve over weeks

To confirm that the RM and PV correlations reflect the formation of spatial engrams specific to the set of contextual cues presented during S1, we conducted an additional series of experiments using different contexts between S1 and S2 following saline administration (Fig. 5a). As shown in Fig. 5b, RM_corr_ values were indeed significantly lower in all genotypes when the contextual cues were changed between S1 and S2 (p-WT: t(32.97) = −4.46, p < 0.0001; α5-i-KO: t(21.75) = −2.37, p=0.027; α5-pyr-KO: t(7.27) = −3.76, p=0.0066). There were no significant differences in this measure between genotypes (genotype x experimental condition; α5-i-KO: t(65.04) = 1.20, p = 0.23; α5-pyr-KO: t(56.82) = −1.52, p = 0.13). Similar statistical conclusions were made with PV_corr_ (Supplementary Fig. 10). These findings support the use of RM and PV correlations as neuronal correlates of spatial memory.

**Fig. 5.**
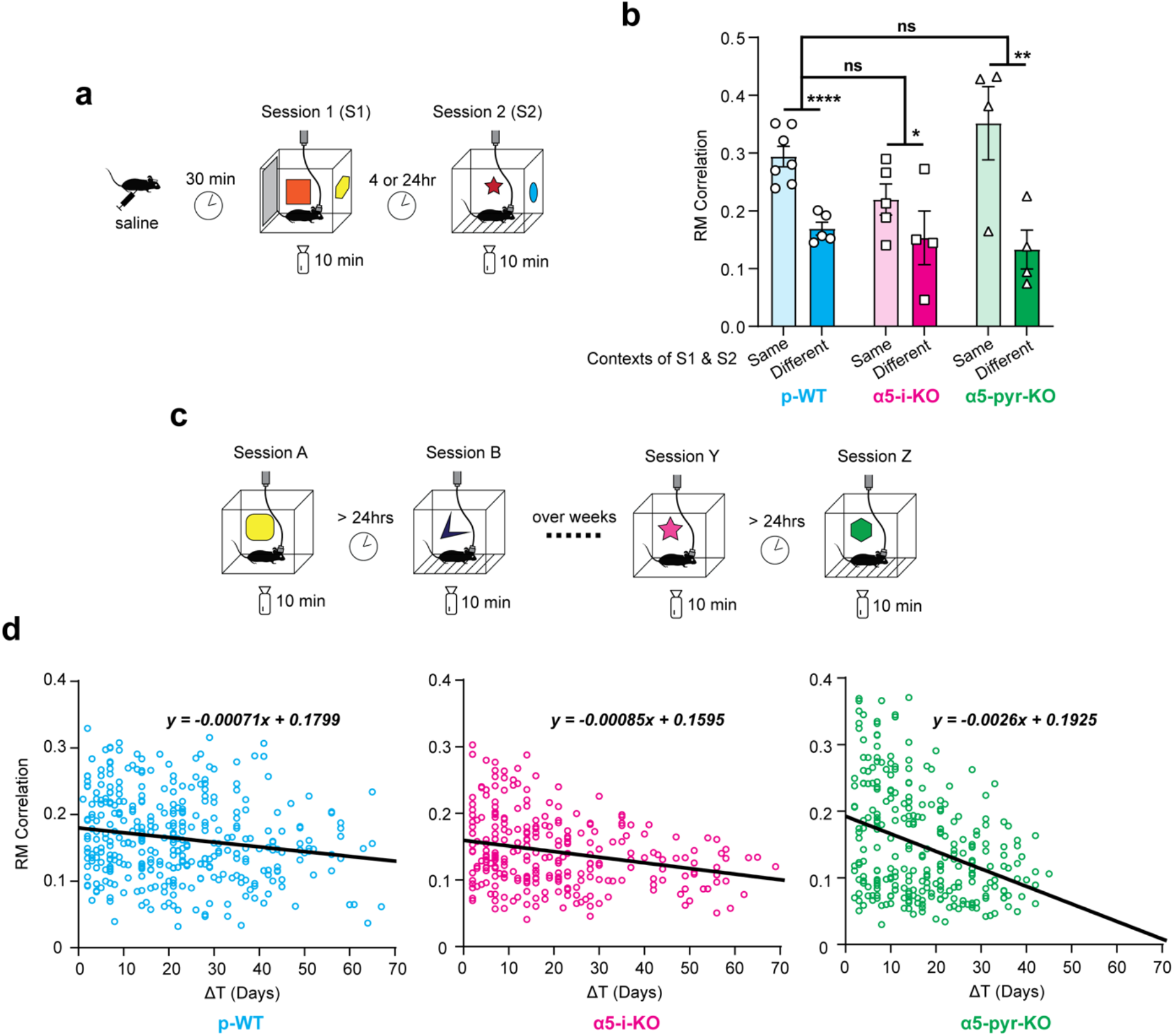
Hippocampal spatial engrams display context specificity. **a** Experimental paradigm used to investigate context specificity of PV and RM correlations as contextual memory correlates. Mice were injected with saline 30 minutes prior to S1, and local contextual cues were changed before S2. **b** Summarized results from different-contexts experiments. Changing contextual cues significantly (and similarly) reduced RM correlations for all three genotypes (**p < 0.01, ****p < 0.0001). **c** Experimental paradigm used to investigate the ‘residual correlation’ observed in different-contexts experiments in part A. Mice were exposed to novel contexts (different local cues) over weeks in the same general environment (identical distal cues). **d** Residual RM correlation ‘drift’ over weeks for all three genotypes with derived linear regression models. ‘Residual correlations’ were higher when two sessions were conducted closer in time, and y-intercepts of the linear regression models closely match results obtained in different-contexts experiments in part A. α5-pyr-KO mice had significantly faster drift and significantly higher ‘residual correlations’ at early time points (y-intercept).

Although both RM and PV correlations were reduced in different-contexts experiments compared to same-context experiments, they still were significantly higher than ‘0’, i.e. the location-shuffled null values (Fig. 4c, dashed lines). These ‘residual correlations’ suggest that some component of the hippocampal activity pattern reflects a more general aspect of the environment (distal cues), such as the room in which experiments were conducted or the plexiglass arena surrounded by a blackout curtain into which the mice were placed, as opposed to the unique sensory cues within the arena that constituted the ‘context’ (local cues) ^64^. Because we had always presented a new set of local cues for every S1 and S2 pair, we were able to conduct analyses analogous to the different-context experiments by treating each S2 as a unique recording session with novel local cues yet identical distal cues, thereby analyzing the ‘residual correlations’ for up to ten weeks (Fig. 5c). We observed that there was a slow ‘drift’ of the ‘residual correlations’ over weeks for all three genotypes, as shown by linear regression models with significantly non-zero slopes (pWT: F(1, 361) = 11.36, p = 0.0008; α5-i-KO: F(1, 272) = 25.31, p < 0.0001; α5-pyr-KO: F(1, 275) = 36.91, p < 0.0001) (Fig. 5d). Moreover, ‘residual correlations’ were greater when two sessions were close in time (e.g. 5 days), as opposed to farther in time (e.g. 60 days). The y-intercepts of these linear models represent predicted mean RM_corr_ of different-contexts experiments conducted close in time, and as expected, the y-intercepts closely matched those we obtained in different-contexts experiments. However, α5-pyr-KO mice had faster ‘drift’ in ‘residual correlations’ compared to p-WT and α5-i-KO mice (95% CI of linear regression model slopes: p-WT = −0.0011 – −0.00029; α5-i-KO = −0.0012 – −0.00051; α5-pyr-KO = −0.0035 – −0.0018). Again, PV correlations provided similar findings (Supplementary Fig. 11). These findings lead us to speculate that spatial engrams are formed by a combination of experimentally altered local cues and unaltered distal cues (i.e. different levels in a hierarchical memory system ^65^), and that the representation of the environment drifts over a time frame corresponding to slow components of system consolidation ^66^.

We conducted a final set of experiments analogous to the same-context (S1-S2 pair) experiments but using a separate group of four WT mice, where pairs of sessions were separated by either 4 hours or 24 hours, to test whether spatial engrams have similar stability over 4 hours vs. 24 hours (Supplementary Fig. 12). RM and PV correlations were essentially identical, indicating that the spatial engrams measured at 4 hours provide a good measure of their strength at 24 hours (and vice versa) – a more typical measure of long-term memory in behavioral studies, including those that we used for CPFE tests of memory (Fig. 2).

## Discussion

Results from both behavioral and Ca^2+^ imaging experiments demonstrated that α5-GABA_A_Rs on interneurons are essential targets for learning suppression by the archetypal GABAergic general anesthetic etomidate. The calcium imaging results further showed that in addition to this pharmacological role, interneuronal α5-GABA_A_Rs play a physiological role in promoting spatial learning. Ca^2+^ imaging experiments provided additional mechanistic insights by showing that etomidate suppresses the initial development of place cells and interferes with the formation and/or stability of a spatial engram encoded by a combination of place cells and non-place cells. Insofar as spatial engrams serve as neural correlates of hippocampal memories, our results from α5-pyr-KO mice, in which etomidate was able to suppress engrams, and α5-i-KO mice, in which it was not, indicate that etomidate’s actions on interneuronal α5-GABA_A_Rs are both necessary and sufficient for this effect. This conclusion challenges the longstanding and widely accepted model of memory modulation occurring exclusively through α5-GABA_A_R-mediated tonic and slow phasic inhibition on pyramidal neurons ^27, 28, 29, 32, 67, 68^.

The role of α5-GABA_A_Rs on pyramidal neurons is less clear. Whereas selective elimination of these receptors did prevent etomidate from suppressing freezing in the CPFE paradigm (Fig. 2), it did not prevent etomidate from suppressing the formation of place cells in a novel environment (Fig. 3), or from interfering with the development of spatial engrams (Fig. 4). In other words, although etomidate prevented α5-pyr-KO mice from forming a hippocampal contextual representation, mice were still able to associate the arena with the footshock. A likely explanation is that mice usually rely upon hippocampus to overcome the immediate shock deficit in CPFE ^44, 45^, but when the hippocampus is damaged or impaired prior to training, mice can form context – shock associations using other brain regions and learning strategies ^69, 70, 71^. Extrahippocampal regions that mediate these alternate learning strategies include medial prefrontal cortex ^72^, retrosplenial cortex ^73, 74, 75^ and dorsolateral striatum ^76^. It is also possible that the neural correlates that we focused on here do not completely capture the behaviorally relevant contributions of hippocampal activity to shock-elicited freezing. Indeed, others have reported that neural correlates in some learning tasks correspond better than others to behavioral readouts, such as discrimination tasks that depend on pattern separation in DG, but for which behaviors are more consistently aligned with CA1 firing ^77^. Also, hippocampal spatial mapping occurs differently when mice navigate in a multisensory environment compared to one that engages only a single sensory modality ^78^, so our use of a multisensory rich cue set in Ca^2+^ experiments compared to only visual cues in CPFE experiments may have influenced our findings. In any case, results from Ca^2+^ imaging studies provided clear evidence that α5-GABA_A_Rs on pyramidal neurons do not mediate the impairment of hippocampal function produced by etomidate.

The methods we used – repeatedly exposing mice to novel sets of contextual cues several times per week for up to twelve weeks – provided robust cellular correlates of spatial memory, and they allowed us to develop full dose-response relationships from groups of only 4-6 mice of each genotype. Not only should this approach be applicable to a wide range of questions, but it also has the added benefits of efficient resource utilization and minimization of animal welfare concerns by reducing the total number of animals used.

Here we have used the term “spatial engram” to refer to the correspondence between the recurrence of spatially modulated neuronal activity patterns (Fig. 4) and the formation or suppression of hippocampus-dependent contextual memories (Fig. 2). In this regard, spatial engrams incorporate characteristics of both place fields as cognitive maps ^47, 79^ and engrams as revealed and captured by the activation of learning-related IEGs and replayed through optogenetic activators ^61, 62, 63^. However, spatial engrams differ from place cell maps and IEG-defined engrams (a collection of engram cells) in several important ways: i) they include both place cells and non-place cells; ii) they presumably include both engram cells and non-engram cells ^59, 80^; and iii) they represent ensemble activity of a large proportion of the population rather than a restricted subset of highly active cells. These spatially modulated firing patterns, which other investigators who have used the same recording methods have also observed ^55, 56, 64, 81^, embody many of the characteristics expected of neuronal correlates of memory. These include the ability to retain stable representations of an environment over extended periods of time ^55, 64^, the ability to discriminate between arenas with different cues but at the same time to generalize between arenas in the same general environment ^64^ (and Fig. 5), and fluctuating activity patterns that create unique episodic time stamps ^82^. Here we add to this collection of attributes the finding that drugs that prevent learning also directly interfere with the development of stable neuronal ensembles – thus extending it from a set of correlations to include an interventional test. Admittedly, these findings do not yet match the extensive evidence developed over the past decade indicating that IEG-captured engram cells represent the physical substrate of memories themselves ^62^. To develop a comparable level of evidence would require reinstating the activity of the entire ensemble in a manner that drives memory-guided behavior, but such a method has not yet been developed. Nevertheless, it is likely that spatial engrams do include IEG-captured engram cells; ecphory might therefore occur, but this point is speculative. Furthermore, just as a set of bona-fide engram cells observed in the hippocampus or elsewhere in the brain does not constitute an entire memory trace but is just one portion of an engram complex ^83^, the spatial engrams that we describe here are likely one component of a memory trace that extends across multiple structures ^84^, to capture the memory of the arena, the general environment, and the context within which the experiments take place.

Which types of interneurons might be responsible for the present findings? It is counterintuitive that suppressing interneuron activity through positive modulation of their α5-GABA_A_Rs would impair memory – at least if one considers inhibitory interneurons only in terms of controlling the depolarization of pyramidal neurons. Reducing this constraining influence would instead lead to net disinhibition, enhancing the ability of pyramidal neurons to generate dendritic bursts, become place cells, and establish networks that store memories ^32, 85, 86^. However, interneurons also target other interneurons, creating positive feedback loops that can support the induction of long-term potentiation and associative learning and memory ^87, 88^. Interrupting these disinhibitory loops might then be the key. In this regard, interneurons that express vasoactive intestinal peptide (VIP+ INs) are an obvious candidate, since they preferentially target other interneurons. However, VIP+ INs were reported to not express α5-GABA_A_Rs in human or mouse prefrontal cortex ^89^, and an examination of the distribution of Gabra5 mRNA in hippocampal CA1 interneurons using a previously published database ^90^ confirms that this is the case in hippocampus as well, as Gabra5 levels are lowest in Vip+ of all interneuron classes (Supplementary Fig. 13). At the other end of the Gabra5 expression spectrum are Cck+ basket cells – their Gabra5 expression levels are the highest of all (Supplementary Fig. 13), they are densely expressed in dorsal CA1 ^91^, and their selective activation has been shown to enhance memory ^92^. Therefore, depression of Cck+ basket cells via α5-GABA_A_Rs is one possible mechanism. Neurogliaform and ivy cells also express Gabra5, at intermediate levels (Supplementary Fig. 13), and they also receive GABA_A_R-mediated slow IPSCs ^93^, which on pyramidal neurons are mediated by α5-GABA_A_Rs ^32, 67, 94^. In addition, they have high-density release sites, and they inhibit other interneurons as well as pyramidal cells via firing patterns that are time-locked to the theta oscillation ^93, 95^. Therefore, these cells may also be involved. Finally, Sst+ INs, which receive α5-GABA_A_R,_slow_ IPSCs ^96^ and play essential roles in the formation and control of contextual fear conditioning ^97, 98^, are also good candidates for the control of learning and memory through interneuronal α5-GABA_A_Rs.

A role for α5-GABA_A_Rs located on interneurons in controlling hippocampus-dependent memory may have wide ramifications. Changes in α5-GABA_A_R expression or activation have been proposed to contribute to memory deficits in Alzheimer’s disease, schizophrenia, autism, aging, and in the early postoperative period ^22, 99, 100, 101, 102, 103^. Even if they are not the pathological basis *per se*, modulation of α5-GABA_A_Rs has also been found effective in preclinical models of depression and Down syndrome ^19, 20, 21^. It should be noted that our findings also support important roles for α5-GABA_A_Rs on pyramidal neurons; indeed, there is evidence that pyramidal neuron α5-GABA_A_Rs specifically regulate certain memory domains ^46^. However, most studies focused on cognition have assumed that effects come from direct modulation of pyramidal neurons; the idea that they might derive instead (or in addition) from modulation of interneurons opens interesting possibilities. For example, modulation of α5-GABA_A_Rs on different cell types might help explain why both α5-PAMs and α5-NAMs show efficacy in treatment of animal models of depression ^18, 19, 21^. The recognition that α5-GABA_A_Rs on interneurons are essential components of the hippocampal memory machinery, together with an emerging understanding of the functional diversity of interneurons, should present new opportunities for discovery and development of targeted therapeutics.

## Methods

### Animals

All experiments were conducted under the oversight and with the approval of the Institutional Animal Care and Use Committee (IACUC) of the University of Wisconsin-Madison. Contextual fear conditioning experiments (CPFE) were conducted on a total of 74 mice from the GAD2-Cre (GAD2^tm2(Cre)Zjh^) x α5-floxed (Gabra5^tm2.1Uru^) line and 90 mice from the CaMKIIa-Cre (Tg(Camk2a-Cre)T29-1Stl) x α5-floxed (Gabra5^tm2.1Uru^) line. Ca^2+^ imaging experiments (GCaMP6f) were conducted on 7 p-WT mice (α5-floxed, Cre-negative), 5 α5-i-KO mice, 4 α5-pyr-KO mice, and 4 wild type C57BL6/J mice. Mice of both sexes were used for CPFE experiments (85 females, 79 males). Only male mice were used for GCaMP6f imaging because their larger size facilitated lens and baseplate implantation.

All mice were generated and maintained on the C57BL/6J background. Pseudo-WT (p-WT) and their interneuron-specific Gabra5 knockout littermates (α5-i-KO and α5-pyr-KO) were created in a three-step process. F0 mice were homozygous α5-floxed (α5:fl/fl) kindly provided by the University of Zurich ^104^, homozygous GAD2-IRES-Cre (JAX strain # 010802) ^105^, hemizygous Viaat-Cre (JAX strain # 017535) ^106^, or homozygous CaMKIIa-Cre (JAX strain # 005359) ^107^ mice obtained from the Jackson Laboratory (Bar Harbor, Maine, USA). As expected, F1 offspring were all heterozygous for the α5-floxed gene (α5:fl/-) and the GAD2-Cre or CaMKIIa-Cre gene, and approximately one-half were also hemizygous for the or Viaat-Cre gene. F2 mice were created by crossing a F1 α5:fl/-;Cre+ mouse and a F1 α5:fl/-;Cre-mouse. F3 mice were created by crossing F2 mice that were homozygous for α5-floxed gene (α5:fl/fl;Cre-) with F2 mice also homozygous for α5-floxed gene plus hemizygous for GAD2-Cre, Viaat-Cre, or CaMKIIa-Cre genes (α5:fl/fl;Cre+). F3 mice consisting of (α5:fl/fl;Cre+) and (α5:fl/fl;Cre-) were used as breeding pairs. Experimental mice came from F3 breeding pairs and subsequent generations produced by the same strategy (α5:fl/fl;Cre+) x (α5:fl/fl;Cre-).

We monitored for germ-line recombination by testing for the presence of Cre recombinase plus the α5-floxed allele, the α5-KO allele, and the α5-WT allele. Any mice that were negative for Cre recombinase but positive for the α5-KO allele were considered to have undergone germ-line recombination. We observed a high rate of germ-line recombination in the Viaat-Cre x α5-floxed line, whether Cre was carried by the male or female (~25% of offspring), but never any germ-line recombination in the GAD2-Cre x α5-floxed or CaMKIIa-Cre x α5-floxed lines, whether Cre was carried by the male or female. Therefore, all experiments were conducted using only mice from the GAD2 and CaMKIIa lines.

### Genotyping

Mice were genotyped from tail samples either in-house using traditional, gel based PCR methods, or sent to Transnetyx (Cordova, TN), which uses a TaqMan-based assay for real-time PCR data. For in-house PCR, primers purchased from IDT (Integrated DNA Technologies, Coralville, IA) were as follows (forward, reverse): Gabra5: TGATGGCACACTTCTCTACACC, CTTTGAAAGCATTTCCCGAAGC; GAD2-Cre: CTAGGCCACAGAATTGAAAGATCT,GTAGGTGGAAATTCTAGCATCATCC; Viaat-Cre: GCGGTCTGGCAGTAAAAACTATC, GTGAAACAGCATTGCTGTCACTT.

### Ex-vivo microscopy

To image fluorescent markers td-Tomato and GFP, coronal slices (100-400 um thickness) were cut from three mice and imaged live in cold, oxygenated ACSF. Hippocampal subfields were imaged on an Olympus FluoView FV1000 upright confocal microscope with a water immersion objective (either a LUMPlanFl/IR 40X N.A.=0.8, or a XLUMPLFL 20X N.A.=1.00). Z-series images (at least 40-50μm in Z) were acquired with 488nm and 561nm laser lines (sequential excitation). Several dozen fluorophore-expressing cells were observed in each volume of interest. From these z-series, the cell bodies of fluorophore-expressing cells (either td-Tomato or GFP) were segmented, and average fluorescence (a.u.) in each color was quantified.

### Tissue sections for IHC

Mice (P60 – P120) were anesthetized by isoflurane inhalation and transcardially perfused with 4% PFA/PBS. Post-fixation, brains were incubated in 4% pFA/PBS for 4 hours followed by an overnight incubation in 30% sucrose/PBS at 4° C. Prior to sectioning, brains were incubated in 4.5 pH Na-Citrate buffer overnight and then underwent microwave heating (900 Watts in 80 mL freshly prepared 6 pH Na-Citrate buffer for 90 seconds). The brains were then coronally sectioned at 30 μm with a vibratome (Leica VT1000s).

### Immunohistochemistry

Slices were washed 2x in PBS for 10 min and then incubated in primary antibody solution (0.2 % Triton X-100, 10% Normal Goat Serum (NGS; Thermo Fisher Scientific Cat# 50197Z), rabbit polyclonal anti-GABRA5 (1:200, Origene Cat# TA338505), guinea pig polyclonal anti-GAD2 (1:200, Synaptic Systems Cat# 198 104) overnight at 4° C in a moist chamber with constant agitation (100 rpm). Slices were then washed 3x in PBS for 15 min followed by a 45-minute incubation in secondary antibody solution (2% NGS, 1:200 Goat anti-Rabbit IgG (H+L) Alexa Fluor^™^ 568 Thermo Fisher Scientific Cat# A-11036, 1:200 Goat anti-Guinea Pig IgG (H+L) Alexa Fluor^™^ 647 Thermo Fisher Scientific Cat# A-21450) at room temperature (RT) with continuous agitation. Slices were then washed 3x for 15 minutes for a final time before mounting on slides with a DAPI mounting medium (Abcam Cat# ab104139). Slides were imaged using a Nikon upright FN1 confocal microscope.

### Western blot analysis

A 1:2 dilution of supernatant of the anti-α5-GABA_A_R NeuroMab clone N415/24 was used. There were three animals per genotype and gender. Single animals from each group were run per gel (3 gels total). Expression levels were quantified by densitometry and normalized to β-actin (42 kDa). The means from the three WT males or three WT females, corrected for β-actin levels, were set to 1.0.

### Behavioral experiments

Behavioral studies were carried out at the Waisman Center Rodent Models Core facility at the University of Wisconsin-Madison. Mice were transferred from the primary animal care unit in which they were bred and raised to the Waisman Center animal care unit at least one week prior to initiating behavioral experiments. Studies of contextual fear conditioning were carried out first, followed by elevated plus maze, and then thermal sensitivity, with 3-4 days between experiments.

### Context preexposure facilitation effect (CPFE)

We used a preexposure-dependent contextual fear conditioning paradigm adapted from Cushman et al., 2012 ^108^. This paradigm, which is often referred to in the literature as the Context Preexposure Facilitation Effect (CPFE) paradigm ^40, 41, 42^ takes advantage of the so-called “immediate shock deficit” ^39, 43^, wherein animals that are shocked immediately (within several seconds) upon entry into a novel environment do not freeze on subsequent re-exposure, whereas mice that had been exposed on a prior day do exhibit a freezing response. The proposed explanation is that preexposed mice establish a hippocampus-dependent representation of the environment (i.e. contextual memory), which takes several minutes, and this pre-formed memory can be recalled rapidly on re-exposure, for subsequent association with the aversive stimulus ^44, 45^. This paradigm was chosen because etomidate, as an anesthetic, might suppress freezing to context during a standard contextual conditioning paradigm, not by preventing the formation of a contextual memory (a hippocampus-dependent process), but rather by reducing the aversive nature of the shock, or some other aspect of its ability to support aversive conditioning. By administering etomidate during the context exposure phase, separated by one day from the shock exposure phase, that interpretive ambiguity is removed.

The CPFE experiments took place over two weeks. During the first week, the mice were habituated and handled in the behavioral testing room for 10 min a day. During the second week, the mice underwent three experimental phases – context preexposure, conditioning, and recall – on three consecutive days. Mice were brought into the behavioral testing room 30 minutes prior to the testing procedure. On Day 1 (*context preexposure*), mice were injected with either saline or etomidate (7 mg/kg IP), placed back in their home cage for 30 minutes, and then placed into the test chamber for 10 minutes. The test chamber was 20 cm x 20 cm x 30 cm high, constructed of clear acrylic with checkered patterned paper covering three of the four walls and a shock grid floor consisting of stainless-steel bars 2 cm apart, diameter 2 mm. On Day 2 (*conditioning*), mice were placed into the same test chamber, and after 15 sec they were administered a single foot-shock (2 sec, 1 mA). The mice remained in the test chamber for an additional 30 sec (47 sec total time) and they were then returned to their home cage. On Day 3 (*recall*), mice were placed in the test chamber for 8 minutes. The amount of time they spent moving was recorded using FreezeFrame^™^ software. The percentage of time they were ‘immobile’ (movement below a pre-defined threshold level) on Day 1 served as a measure of drug-induced sedation; the percentage of time that they were immobile during the first three minutes of the drug-free test on experimental Day 3 (‘freezing behavior’) served as a quantitative measure of fear memory.

No significant differences between sexes were seen for either Day 1 (sedation) or Day 3 (freezing) for any groups (matching genotype and drug condition, unpaired t-test without correction for multiple comparisons), or in interaction between genotype (p-WT vs KO) and drug condition (saline vs ETOM) for either α5-i-KO or α5-pyr-KO lines (2-way ANOVA), so data from males and females were combined.

### Elevated plus maze (EPM) test

Mice were placed individually in a plus-shaped maze composed of two “open” arms without walls (30 cm L X 5 cm W) and two closed arms (30 cm L X 5 cm W) enclosed by walls (10 cm H) arranged around a center zone (5 cm L X 5 cm W). Over the course of five minutes, the amount of time they spent on the arms and the number of entries to each arm were manually recorded by an experimenter blind to the genotype of the mice. The percent time the animal spent on the open arms and the number of entries to each arm served as a measure of anxiety-like behavior, with more entries and more time spent on the open arms indicating less anxiety.

### Hotplate test

An electronically controlled hotplate (30 cm L X 30 cm W) heated to 55 °C was used to measure sensitivity to a noxious thermal stimulus. The hot plate was turned on 30 minutes prior to testing to ensure that the desired temperature was reached. Mice were then placed individually on the hot-plate and the latency to elicit a nocifensive behavior (e.g., hind paw withdrawal or licking) was manually recorded by the experimenter blinded to the genotype of the mice.

### Ca^2+^ imaging experiments

Twenty male mice (7 p-WT, 5 α5-i-KO, 4 α5-pyr-KO, 4 C57BL/6J) each underwent two stereotaxic surgeries separated by 2-3 weeks. For both procedures, mice were anesthetized with isoflurane ~2% adjusted to maintain immobility and spontaneous ventilation and warmed by block heater to maintain normothermia. Postoperatively mice received a subcutaneous injection of carprofen 5 mg/kg. For the first surgery, a small craniotomy window was made over the right dorsal hippocampus (from bregma: AP = −2.0, ML = −1.6) and 500 nL of virus carrying the genetically encoded calcium indicator GCaMP6f driven by the CaMKIIa promoter (Inscopix Ready-to-Image AAV1-CaMKIIa-GCaMP6f) was injected at DV = −1.6 at a rate of 80 nL/min using a NanoFil syringe with a 35 g needle driven by a UMP3 microsyringe pump and SMARTouch controller (WPI). The needle was kept in place for 2 min, then retracted 200 um, where it remained for an additional 8 minutes before removing slowly. The skin was sutured closed, and the mouse was observed during recovery from surgery and anesthesia then returned to its home cage. For the second surgery, which took place 2-3 weeks later, a larger craniotomy was made at AP = −2.2, ML = −2.1, and the cortex corpus callosum overlaying the hippocampus was aspirated using a 30g blunt needle and cold saline irrigation, taking care not to aspirate the alveus. Once bleeding was controlled with Gelfoam and cold saline irrigation, an integrated GRIN lens (1 mm diameter x 4 mm length) and baseplate were inserted slowly at a 9-degree angle to the midline, until the center of the surface of the lens rested DV = −1.2. The baseplate was cemented in place using Metabond, the surrounding skin was sutured closed, and the mouse was allowed to recover from surgery and anesthesia, then returned to its home cage.

Two to three weeks later, a miniature epifluorescence microscope (Inscopix nVoke) was affixed to the baseplate, and the mouse was placed in the behavioral arena (a 40 cm x 40 cm x 30 cm tall acrylic enclosure) surrounded by blackout curtain, with exam table paper placed on the floor of the arena and allowed to explore freely for 10 minutes. A commutator (Inscopix) was used to prevent tangling of the wire attached to the microscope. During this initial ‘screening session’, the focal plane, illumination, and gain were adjusted to identify the optimal settings to observe cellular activity. If the number of cells with sufficient activity levels was too low (fewer than 60 cells with >5 Ca^2+^ events) the mouse was returned to its home cage and tested one week later. Once sufficient activity was observed, which typically occurred 5-6 weeks following the initial surgery with virus injection, the same settings were used for all subsequent recording sessions, except that the illumination was occasionally adjusted down or up to maintain appropriate fluorescent signal levels.

On each experimental day, two recording sessions separated by either 4 or 24 hours (Session 1 and 2, 10 min each) took place in the same behavioral arena used for the initial screening sessions. For Session 1, the arena contained a unique set of visual, olfactory, and tactile cues forming a ‘novel context’. Visual cues consisted of 8” x 11” sheets of paper mounted onto the outsides of all four walls, with different solid colors, shapes (stars, circles, squares, triangles) and patterns (stripes, swirls, lines) on the four sheets. Olfactory cues consisted of 1 μL of odorant (benzaldehyde, hexanal, alpha-pinene, heptaldehyde, eugenol, or eucalyptol) on filter paper placed within a 35 mm covered culture dish and set in one corner of the arena. Tactile cues consisted of ¼” thick acrylic squares covering one quarter of the arena floor, or various sizes of shallow glass petri dish lids placed either open side up or down, in addition to a sheet of exam table paper covering the floor of the entire arena. Prior to Session 1, a mouse received either saline (control) or etomidate (2-8 mg/kg) by intraperitoneal injection, carried out in a separate room. Thirty minutes after the injection the mouse was brought into the recording room, the miniature microscope was attached to the baseplate, the mouse was placed in the arena, and the light in the arena was turned down to ~2 lux to encourage exploration. A bank of infrared lights above the mouse (12 lux) was used to illuminate the arena so that the mouse could be tracked using an IR-sensitive camera (Basler aca1300-60 gm) mounted below the arena (Fig. 3a and 4A). The video camera and epifluorescent microscopy acquisition were synchronized using a hardware trigger controlled by Ethovision software (Noldus Information Technology). At the end of the 10-min recording session, the miniature microscope was detached, and the mouse was returned to its home cage in a separate room. For Session 2, the mouse was brought back into the recording room, the miniature microscope was mounted to the baseplate, and the mouse was placed back into the arena containing the same set of sensory cues used for Session 1.

Experiments using a novel set of cues for Session 1 and unchanged cues for Session 2 (same-context experiments) took place 2-3 times per week, with at least one day between two consecutive pairs of recording sessions. For different-contexts experiments, altered sets of sensory cues were used for Session 1 vs. Session 2. For each mouse, experiments continued for up to 10 weeks (16 weeks post-virus injection), at which time the number of cells with sufficient activity was typically declining.

Behavioral tracking recordings were analyzed using Noldus EthoVision XT15 and exported as csv files. These csv files summarized the mice’s XY-location (body center) as a function of time. All behavioral tracking csv files were organized into a specific data structure with experimental time, animal ID, genotype, and related experimental conditions. The in-vivo calcium imaging recordings were also organized into a matching data structure. An excel metadata sheet for all animals and all experiments allows for programmatic reference (MATLAB 2021a).

### Initial processing of calcium imaging recordings

We utilized the MATLAB API package embedded within Inscopix Data Processing Software v1.8.0 (IDPS v1.8.0) for this initial stage of analysis. To begin, each raw recording was ‘preprocessed’ to remove artifacts, then both spatially and temporally downsampled by a factor of two to improve processing speed without compromising the resolution of cellular calcium dynamics. These recordings were then spatially filtered (parameters: low cutoff = 0.005 pixel^−1^; high cutoff = 0.5 pixel^−1^) to remove low and high spatial frequency content for better contrast and smoother frames, and frames were motion-corrected based on an arbitrarily chosen region of interest (ROI). To improve visualization of calcium events and prepare for the second-stage analysis by CNMF-E algorithm, a ΔF/F transformation was applied and a maximum projected image of the entire field of view was acquired.

### Extraction of cellular activities and longitudinal registration of cell maps

The constrained nonnegative matrix factorization for microendoscopic data (CNMF-E) analysis was applied to the recordings after the initial processing stage, with analysis parameters optimized according to published methods ^109^ and visual inspections. Noisy calcium traces of detected cells were deconvolved using the ‘online active set method to infer spikes’ (OASIS) method ^110^. Once deconvolved, calcium events were inferred from calcium traces based on the biokinetics of GCaMP6f, using a threshold of 4 median absolute deviations (MAD) ^111^. Several analyzed cellsets were randomly chosen to verify that inferred spikes accurately reflected calcium signals from detected cells, based on visual inspection. To exclude non-cellular activities picked up by the CNMF-E algorithm, we required that detected cells must have exactly one spatial component, appropriate size, and more than 5 detected calcium events over a 10-minute recording session. Finally, cellsets of all recordings obtained from an individual mouse were longitudinally registered, so that the activity of each unique cell could be traced between and across recording sessions ^57^. These data were then exported as csv files that included global cell IDs, the timestamps (beginning of rising phase) of inferred calcium events, and the fluorescence amplitude of calcium events in MAD.

### Place cell and spatial engram analysis

For the second stage of analysis, we used custom-written MATLAB functions to merge behavioral tracking and calcium event data to characterize place cells and spatial engrams (Supplementary Fig. 3). Behavioral data were automatically analyzed and exported from Noldus EthoVision XT15 software. *Behavioral Tracking Analysis*: Mouse speed as a function of time was derived from its XY-location (mouse body center) as a function of time (resolution of 0.04 sec) exported from Noldus software and smoothed with a triangular window (size = 0.3 sec). Mobility was then quantified as the fraction of time that the mouse spent actively exploring (speed > 1 cm/sec) over a 10-minute recording session. Next, an occupancy-time map was created by dividing the arena into a 15-by-15 matrix (i.e. 225 pixels) and computing the duration (in sec) that a mouse spent within each pixel. *Place Cell Analysis*: This stage of analysis is conceptually shown in Supplementary Fig. 3A. First, calcium event timestamps were matched with behavioral tracking timestamps to find the spatial location of the mouse at the beginning of rising phase of each calcium event. Second, calcium events that occurred during periods of inactivity (speed < 1cm/sec) were excluded. Third, for each cell, an amplitude-weighted *calcium event map* was created by duplicating calcium events according to their amplitudes (~1-8 MAD) and distributing them along the track of the mouse, at intervals of 40 msec, over the rising phase of the GCaMP6f fluorescence signal. Fourth, for each cell, a *calcium event rate map* (events/sec; 15-by-15 matrix) was created by dividing the calcium event map by the occupancy-time map, pixel-by-pixel. Fifth, for each cell, the mutual information (MI) between the calcium event rate map and occupancy-time map for each cell was calculated according to the formulae:

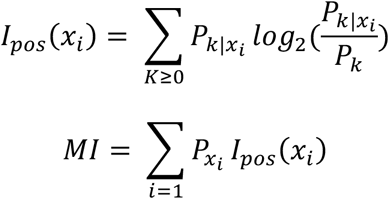

Where:

- *x*_*i*_ refers to one pixel in the binned arena (in our analysis, the arena is depicted by a 15-by-15 matrix).
- *I*_*pos*_(*x*_*i*_) is the positional information (in bits) of pixel *x*_*i*_.
- 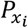 is the probability that the mouse occupies pixel *x*_*i*_, calculated as 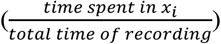
- *P*_*k*_ is the probability of observing *k* calcium events
- 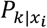 is the conditional probability of observing *k* calcium events in pixel *x*_*i*_, which is equivalent to calcium event rate in pixel *x*_*i*_.
- *MI* is the mutual information (in bits per second) between calcium event rate and mouse’s spatial location.

(adapted from: ^64, 112^. Finally, for each cell, a null distribution of time-shuffled MI values was created by 1000 random circular permutations of calcium event timestamps; this null distribution was used to calculate p(MI). To facilitate visualization of place specific firing, rate maps in figures are presented as 50×50 color-coded matrices. *Rate Map (RM) Correlation Analysis*: This analysis is conceptually shown in Supplementary Fig. 3B. First, for each cell, calcium event rate maps were smoothed by a Gaussian filter (radius = 10 cm, sigma = 6 cm) to generate smoothed calcium event rate maps. Second, for each cell, Pearson’s correlation coefficient (PCC) was calculated between the vectorized smoothed calcium event rate maps from S1 and S2 (or more broadly, between any two arbitrarily chosen sessions). Third, to detect and account for possible coherent rotation of cells’ event rate maps ^64^, we calculated RM correlation by rotating maps from one session only (either S1 or S2) by 0°, 90°, 180°, and 270°, and the rotation that resulted in maximum mean(RM) was designated as angle of coherent rotation—which usually was 0°. Fourth, for each pair of sessions, the distribution of PCCs from all coactive cells (i.e. those with >5 calcium events during both sessions) was referred to as RM correlation distribution. Finally, for each pair of sessions, a null distribution of RM correlation values was created by 100 random spatial permutations of smoothed calcium rate maps. *Population Vector (PV) Correlation Analysis*: This analysis is conceptually shown in Supplementary Fig. 3C. First, for each pair of sessions, smoothed calcium event rate maps of all coactive cells were organized into two 3-dimensional matrices, thus forming for each pixel a pair of population vectors (PVs). Second, for each pixel, the PCC between the two PVs was calculated, resulting in a PV correlation distribution. Third, to detect and account for coherent rotation, we calculated PV correlation by using the same rotation procedure as in RM correlation computation, and again an angle of coherent rotation based on PV is determined—which again usually was 0° (though RM and PV did not always agree). Fourth, for each pair of sessions, a null distribution of PV correlation values was created by 100 random permutations of PVs for all pixels.

#### Data presentation and statistical analysis

Statistical comparisons for behavioral studies (CPFE, elevated plus maze, and hot plate) were made using 2-way ANOVA followed by Šídák’s multiple comparisons test, where interaction factors (e.g. drug and genotype) are reported as ‘A x B’ (Graphpad Prism v. 9.2.0). Comparisons for Ca^2+^ imaging data were made by fitting linear mixed effects models using the ‘lmer’ package, with statistical significance derived from Kenward-Roger type III & likelihood ratio tests ^113^, implemented in RStudio, 2021.09.0 Build 351 (R Core Team (2021). R: A language and environment for statistical computing. R Foundation for Statistical Computing, Vienna, Austria. URL https://www.R-project.org/). To evaluate place cell proportions in single sessions, we used a model with drug (saline or etomidate dose) and genotypes as fixed effects, and mouse ID and experimental date as random effects. To evaluate RM_corr_ and PV_corr_ from pairs of sessions we used a model with drug and genotypes as fixed effects, and cell ID (RM_corr_ only), mouse ID, and date as random effects.

## Materials Availability

All materials used in the analysis will be made available upon reasonable request from the Lead Contact.

## Data and Code Availability

The data and analysis code generated in this study are available upon reasonable request to the corresponding authors.

## Acknowledgements

We thank the University of Zurich for providing the floxed Gabra5 mice under Material Transfer Agreement.

## Funding

National Institutes of Health grant R01 GM118801 (RAP)

National Institutes of Health grant 1R01GM128183 (UR and RAP)

National Institutes of Health grant U54HD090256 (UW-Madison, Waisman Center Core)

National Institutes of Health grant 1S10OD025040-01 (UW-Madison, Shared Instrumentation Facility)

National Institutes of Health grants MH061876 and NS097362 (ERC)

National Institutes of Health grant RO1-MH62122 (MSF)

E.R.C. is an Investigator of the Howard Hughes Medical Institute

## Author contributions

Conceptualization: RAP

Methodology: MZ, AA, MGP, CChu, BMK, CCasey, RL, DR, HH, ERC, RN, UR, MSF, RAP

Investigation: MZ, AA, MGP, RL, DR, RN, RAP

Funding acquisition: ERC, UR, MSF, RAP

Supervision: ERC, UR, MSF, RAP

Writing – original draft: MZ, RAP

Writing – review & editing: MZ, AA, MGP, CChu, BMK, CCasey, RL, DR, HH, ERC, RN, UR, MSF, RAP

## References

1. Olsen RW. GABA(A) receptor: Positive and negative allosteric modulators. Neuropharmacology 136, 10–22 (2018).

2. Engin E, Benham RS, Rudolph U. An Emerging Circuit Pharmacology of GABAA Receptors. Trends Pharmacol Sci 39, 710–732 (2018).

3. Ghoneim MM, Block RI. Learning and memory during general anesthesia: an update. [Review] [152 refs]. Anesthesiology 87, 387–410 (1997).

4. Sandin RH, Enlund G, Samuelsson P, Lennmarken C. Awareness during anaesthesia: a prospective case study. The Lancet 355, 707–711 (2000).

5. Mashour GA, et al. Prevention of intraoperative awareness with explicit recall in an unselected surgical population: a randomized comparative effectiveness trial. Anesthesiology 117, 717–725 (2012).

6. Osterman JE, Hopper J, Heran WJ, Keane TM, van der Kolk BA. Awareness under anesthesia and the development of posttraumatic stress disorder. Gen Hosp Psychiatry 23, 198–204 (2001).

7. Benkwitz C, Liao M, Laster MJ, Sonner JM, Eger EI, 2nd, Pearce RA. Determination of the EC50 amnesic concentration of etomidate and its diffusion profile in brain tissue: implications for in vitro studies. Anesthesiology 106, 114–123 (2007).

8. Forman SA. Clinical and molecular pharmacology of etomidate. Anesthesiology 114, 695–707 (2011).

9. Cheng VY, et al. Alpha5GABAA receptors mediate the amnestic but not sedative-hypnotic effects of the general anesthetic etomidate. J Neurosci 26, 3713–3720 (2006).

10. Martin LJ, Oh GH, Orser BA. Etomidate targets alpha5 gamma-aminobutyric acid subtype A receptors to regulate synaptic plasticity and memory blockade. Anesthesiology 111, 1025–1035 (2009).

11. Sur C, Quirk K, Dewar D, Atack J, McKernan R. Rat and human hippocampal alpha5 subunit-containing gamma-aminobutyric AcidA receptors have alpha5 beta3 gamma2 pharmacological characteristics. Mol Pharmacol 54, 928–933 (1998).

12. Sur C, Fresu L, Howell O, McKernan RM, Atack JR. Autoradiographic localization of alpha5 subunit-containing GABAA receptors in rat brain. Brain Res 822, 265–270 (1999).

13. Tulving E, Markowitsch HJ. Episodic and declarative memory: role of the hippocampus. Hippocampus 8, 198–204 (1998).

14. Chambers MS, et al. Identification of a novel, selective GABA(A) alpha5 receptor inverse agonist which enhances cognition. JMedChem 46, 2227–2240 (2003).

15. Chambers MS, et al. An orally bioavailable, functionally selective inverse agonist at the benzodiazepine site of GABAA alpha5 receptors with cognition enhancing properties. JMedChem 47, 5829–5832 (2004).

16. Atack JR. Preclinical and clinical pharmacology of the GABAA receptor alpha5 subtype-selective inverse agonist alpha5IA. Pharmacol Ther 125, 11–26 (2010).

17. Santrac A, et al. Effect of MP-III-022, positive modulator of alpha 5 containing GABAA receptors, on learning and memory. In: European Neuropsychopharmacology) (2019).

18. Prevot TD, et al. Novel Benzodiazepine-Like Ligands with Various Anxiolytic, Antidepressant, or Pro-Cognitive Profiles. Mol Neuropsychiatry 5, 84–97 (2019).

19. Fischell J, Van Dyke AM, Kvarta MD, LeGates TA, Thompson SM. Rapid Antidepressant Action and Restoration of Excitatory Synaptic Strength After Chronic Stress by Negative Modulators of Alpha5-Containing GABAA Receptors. Neuropsychopharmacology 40, 2499–2509 (2015).

20. Martinez-Cue C, et al. Reducing GABAA alpha5 Receptor-Mediated Inhibition Rescues Functional and Neuromorphological Deficits in a Mouse Model of Down Syndrome. J Neurosci 33, 3953–3966 (2013).

21. Zanos P, et al. A Negative Allosteric Modulator for alpha5 Subunit-Containing GABA Receptors Exerts a Rapid and Persistent Antidepressant-Like Action without the Side Effects of the NMDA Receptor Antagonist Ketamine in Mice. eNeuro 4, (2017).

22. Zurek AA, et al. α5GABAA receptor deficiency causes autism-like behaviors. Ann Clin Transl Neurol 3, 392–398 (2016).

23. Prevot T, Sibille E. Altered GABA-mediated information processing and cognitive dysfunctions in depression and other brain disorders. Mol Psychiatry, (2020).

24. Dawson GR, et al. An inverse agonist selective for alpha5 subunit-containing GABAA receptors enhances cognition. J Pharmacol Exp Ther 316, 1335–1345 (2006).

25. Gill KM, Grace AA. The role of alpha5 GABAA receptor agonists in the treatment of cognitive deficits in schizophrenia. Curr Pharm Des 20, 5069–5076 (2014).

26. Maramai S, Benchekroun M, Ward SE, Atack JR. Subtype Selective gamma-Aminobutyric Acid Type A Receptor (GABA(A)R) Modulators Acting at the Benzodiazepine Binding Site: An Update. J Med Chem 63, 3425–3446 (2020).

27. Wang DS, Orser BA. Inhibition of learning and memory by general anesthetics. Can J Anaesth 58, 167–177 (2011).

28. Orser BA. Lifting the fog around anesthesia. Sci Am 296, 54–61 (2007).

29. Hemmings HC, et al. Towards a Comprehensive Understanding of Anesthetic Mechanisms of Action: A Decade of Discovery. Trends Pharmacol Sci 40, 464–481 (2019).

30. Sperk G, Schwarzer C, Tsunashima K, Fuchs K, Sieghart W. GABA(A) receptor subunits in the rat hippocampus I: immunocytochemical distribution of 13 subunits. Neuroscience 80, 987–1000 (1997).

31. Caraiscos VB, et al. Tonic inhibition in mouse hippocampal CA1 pyramidal neurons is mediated by {alpha}5 subunit-containing {gamma}-aminobutyric acid type A receptors. ProcNatlAcadSciUSA 101, 3662–3667 (2004).

32. Schulz JM, Knoflach F, Hernandez MC, Bischofberger J. Dendrite-targeting interneurons control synaptic NMDA-receptor activation via nonlinear alpha5-GABAA receptors. Nature communications 9, 3576 (2018).

33. Salesse C, Mueller CL, Chamberland S, Topolnik L. Age-dependent remodelling of inhibitory synapses onto hippocampal CA1 oriens-lacunosum moleculare interneurons. J Physiol 589, 4885–4901 (2011).

34. Petrache AL, et al. Selective Modulation of α5 GABA(A) Receptors Exacerbates Aberrant Inhibition at Key Hippocampal Neuronal Circuits in APP Mouse Model of Alzheimer’s Disease. Frontiers in cellular neuroscience 14, 568194 (2020).

35. Rodgers FC, et al. Etomidate Impairs Long-Term Potentiation In Vitro by Targeting alpha5-Subunit Containing GABAA Receptors on Nonpyramidal Cells. J Neurosci 35, 9707–9716 (2015).

36. Zarnowska ED, et al. Etomidate blocks LTP and impairs learning but does not enhance tonic inhibition in mice carrying the N265M point mutation in the beta3 subunit of the GABA(A) receptor. Neuropharmacology 93, 171–178 (2015).

37. Figueroa AG, Benkwitz C, Surges G, Kunz N, Homanics GE, Pearce RA. Hippocampal β2-GABA(A) receptors mediate LTP suppression by etomidate and contribute to long-lasting feedback but not feedforward inhibition of pyramidal neurons. J Neurophysiol 126, 1090–1100 (2021).

38. Mederos S, Perea G. GABAergic-astrocyte signaling: A refinement of inhibitory brain networks. Glia 67, 1842–1851 (2019).

39. Yavas E, Trott JM, Fanselow MS. Sexually dimorphic muscarinic acetylcholine receptor modulation of contextual fear learning in the dentate gyrus. Neurobiol Learn Mem 185, 107528 (2021).

40. Rudy JW. Context representations, context functions, and the parahippocampal-hippocampal system. Learn Mem 16, 573–585 (2009).

41. Miller LA, Heroux NA, Stanton ME. NMDA receptors and the ontogeny of post-shock and retention freezing during contextual fear conditioning. Dev Psychobiol 62, 380–385 (2020).

42. Pinizzotto CC, Heroux NA, Horgan CJ, Stanton ME. Role of dorsal and ventral hippocampal muscarinic receptor activity in acquisition and retention of contextual fear conditioning. Behav Neurosci 134, 460–470 (2020).

43. Fanselow MS. Factors governing one-trial contextual conditioning. Anim Learn Behav 18, 364–270 (1990).

44. Fanselow MS. Contextual fear, gestalt memories, and the hippocampus. Behav Brain Res 110, 73–81 (2000).

45. Krasne FB, Cushman JD, Fanselow MS. A Bayesian context fear learning algorithm/automaton. Frontiers in behavioral neuroscience 9, 112 (2015).

46. Engin E, Sigal M, Benke D, Zeller A, Rudolph U. Bidirectional regulation of distinct memory domains by alpha 5-subunit-containing GABA(A) receptors in CA1 pyramidal neurons. Learn Mem 27, 423–428 (2020).

47. O’Keefe J, Nadel L. The hippocampus as a cognitive map. Clarendon Press (1978).

48. Grieves RM, Jeffery KJ. The representation of space in the brain. Behav Processes 135, 113–131 (2017).

49. O’Keefe J, Dostrovsky J. The hippocampus as a spatial map. Preliminary evidence from unit activity in the freely-moving rat. Brain Res 34, 171–175 (1971).

50. Wilson MA, McNaughton BL. Dynamics of the hippocampal ensemble code for space. Science 261, 1055–1058 (1993).

51. Kentros C, Hargreaves E, Hawkins RD, Kandel ER, Shapiro M, Muller RV. Abolition of long-term stability of new hippocampal place cell maps by NMDA receptor blockade. Science 280, 2121–2126 (1998).

52. Thompson LT, Best PJ. Long-term stability of the place-field activity of single units recorded from the dorsal hippocampuus of freely behaving rats. Brain Res 509, 299–308 (1990).

53. Muller RU, Kubie JL. The effects of changes in the environment on the spatial firing of hippocampal complex-spike cells. J Neurosci 7, 1951–1968 (1987).

54. Anderson MI, Jeffery KJ. Heterogeneous modulation of place cell firing by changes in context. J Neurosci 23, 8827–8835 (2003).

55. Ziv Y, et al. Long-term dynamics of CA1 hippocampal place codes. Nat Neurosci 16, 264–266 (2013).

56. Sheintuch L, Geva N, Baumer H, Rechavi Y, Rubin A, Ziv Y. Multiple Maps of the Same Spatial Context Can Stably Coexist in the Mouse Hippocampus. Curr Biol 30, 1467–1476 e1466 (2020).

57. Sheintuch L, et al. Tracking the Same Neurons across Multiple Days in Ca(2+) Imaging Data. Cell Rep 21, 1102–1115 (2017).

58. Rubin A, et al. Revealing neural correlates of behavior without behavioral measurements. Nature communications 10, 4745 (2019).

59. Tanaka KZ, He H, Tomar A, Niisato K, Huang AJY, McHugh TJ. The hippocampal engram maps experience but not place. Science 361, 392–397 (2018).

60. Stefanini F, et al. A Distributed Neural Code in the Dentate Gyrus and in CA1. Neuron 107, 703–716 e704 (2020).

61. Josselyn SA, Kohler S, Frankland PW. Finding the engram. Nat Rev Neurosci 16, 521–534 (2015).

62. Josselyn SA, Tonegawa S. Memory engrams: Recalling the past and imagining the future. Science 367, (2020).

63. Sun X, et al. Functionally Distinct Neuronal Ensembles within the Memory Engram. Cell 181, 410–423.e417 (2020).

64. Kinsky NR, Sullivan DW, Mau W, Hasselmo ME, Eichenbaum HB. Hippocampal Place Fields Maintain a Coherent and Flexible Map across Long Timescales. Curr Biol 28, 3578–3588 e3576 (2018).

65. Bullmore E, Sporns O. Complex brain networks: graph theoretical analysis of structural and functional systems. Nat Rev Neurosci 10, 186–198 (2009).

66. Kim JJ, Fanselow MS. Modality-specific retrograde amnesia of fear. Science 256, 675–677 (1992).

67. Zarnowska ED, Keist R, Rudolph U, Pearce RA. GABAA receptor alpha5 subunits contribute to GABAA,slow synaptic inhibition in mouse hippocampus. J Neurophysiol 101, 1179–1191 (2009).

68. Ali AB, Thomson AM. Synaptic alpha 5 subunit-containing GABAA receptors mediate IPSPs elicited by dendrite-preferring cells in rat neocortex. CerebCortex 18, 1260–1271 (2008).

69. Fanselow MS. From contextual fear to a dynamic view of memory systems. Trends Cogn Sci 14, 7–15 (2010).

70. Wiltgen BJ, Sanders MJ, Anagnostaras SG, Sage JR, Fanselow MS. Context fear learning in the absence of the hippocampus. J Neurosci 26, 5484–5491 (2006).

71. Schimanski LA, Lipa P, Barnes CA. Tracking the course of hippocampal representations during learning: when is the map required? J Neurosci 33, 3094–3106 (2013).

72. Zelikowsky M, et al. Prefrontal microcircuit underlies contextual learning after hippocampal loss. Proc Natl Acad Sci U S A 110, 9938–9943 (2013).

73. Milczarek MM, Vann SD, Sengpiel F. Spatial Memory Engram in the Mouse Retrosplenial Cortex. Curr Biol 28, 1975–1980 e1976 (2018).

74. Coelho CAO, Ferreira TL, Kramer-Soares JC, Sato JR, Oliveira MGM. Network supporting contextual fear learning after dorsal hippocampal damage has increased dependence on retrosplenial cortex. PLoS Comput Biol 14, e1006207 (2018).

75. Czajkowski R, et al. Encoding and storage of spatial information in the retrosplenial cortex. Proc Natl Acad Sci U S A 111, 8661–8666 (2014).

76. Geerts JP, Chersi F, Stachenfeld KL, Burgess N. A general model of hippocampal and dorsal striatal learning and decision making. Proc Natl Acad Sci U S A 117, 31427–31437 (2020).

77. Allegra M, Posani L, Gomez-Ocadiz R, Schmidt-Hieber C. Differential Relation between Neuronal and Behavioral Discrimination during Hippocampal Memory Encoding. Neuron 108, 1103–1112 e1106 (2020).

78. Radvansky BA, Oh JY, Climer JR, Dombeck DA. Behavior determines the hippocampal spatial mapping of a multisensory environment. Cell Rep 36, 109444 (2021).

79. Robinson NTM, et al. Targeted Activation of Hippocampal Place Cells Drives Memory-Guided Spatial Behavior. Cell 183, 1586–1599 e1510 (2020).

80. Ghandour K, et al. Orchestrated ensemble activities constitute a hippocampal memory engram. Nature communications 10, 2637 (2019).

81. Zhu M, Perkins MG, Lennertz R, Abdulzahir A, Pearce RA. Dose-dependent suppression of hippocampal contextual memory formation, place cells, and spatial engrams by the NMDAR antagonist (R)-CPP. Neuropharmacology 218, 109215 (2022).

82. Rubin A, Geva N, Sheintuch L, Ziv Y. Hippocampal ensemble dynamics timestamp events in long-term memory. Elife 4, (2015).

83. Tonegawa S, Liu X, Ramirez S, Redondo R. Memory Engram Cells Have Come of Age. Neuron 87, 918–931 (2015).

84. Hunsaker MR, Kesner RP. Unfolding the cognitive map: The role of hippocampal and extra-hippocampal substrates based on a systems analysis of spatial processing. Neurobiol Learn Mem 147, 90–119 (2018).

85. Bittner KC, Milstein AD, Grienberger C, Romani S, Magee JC. Behavioral time scale synaptic plasticity underlies CA1 place fields. Science 357, 1033–1036 (2017).

86. Geiller T, et al. Local circuit amplification of spatial selectivity in the hippocampus. Nature, (2021).

87. Letzkus JJ, Wolff SB, Luthi A. Disinhibition, a Circuit Mechanism for Associative Learning and Memory. Neuron 88, 264–276 (2015).

88. Leao RN, et al. OLM interneurons differentially modulate CA3 and entorhinal inputs to hippocampal CA1 neurons. Nat Neurosci 15, 1524–1530 (2012).

89. Hu X, Rocco BR, Fee C, Sibille E. Cell Type-Specific Gene Expression of Alpha 5 Subunit-Containing Gamma-Aminobutyric Acid Subtype A Receptors in Human and Mouse Frontal Cortex. Mol Neuropsychiatry 4, 204–215 (2019).

90. Harris KD, et al. Classes and continua of hippocampal CA1 inhibitory neurons revealed by single-cell transcriptomics. PLoS Biol 16, e2006387 (2018).

91. Whissell PD, Cajanding JD, Fogel N, Kim JC. Comparative density of CCK- and PV-GABA cells within the cortex and hippocampus. Front Neuroanat 9, 124 (2015).

92. Whissell PD, et al. Selective Activation of Cholecystokinin-Expressing GABA (CCK-GABA) Neurons Enhances Memory and Cognition. eNeuro 6, (2019).

93. Capogna M, Pearce RA. GABAA, slow: causes and consequences. Trends Neurosci 34, 101–112 (2011).

94. Prenosil GA, Schneider Gasser EM, Rudolph U, Keist R, Fritschy JM, Vogt KE. Specific subtypes of GABAA receptors mediate phasic and tonic forms of inhibition in hippocampal pyramidal neurons. J Neurophysiol 96, 846–857 (2006).

95. Olah S, et al. Output of neurogliaform cells to various neuron types in the human and rat cerebral cortex. Front Neural Circuits 1, 4 (2007).

96. Magnin E, et al. Input-Specific Synaptic Location and Function of the alpha5 GABAA Receptor Subunit in the Mouse CA1 Hippocampal Neurons. J Neurosci 39, 788–801 (2019).

97. Lovett-Barron M, et al. Dendritic inhibition in the hippocampus supports fear learning. Science 343, 857–863 (2014).

98. Szonyi A, et al. Brainstem nucleus incertus controls contextual memory formation. Science 364, (2019).

99. Hauser J, Rudolph U, Keist R, Mohler H, Feldon J, Yee BK. Hippocampal alpha5 subunit-containing GABAA receptors modulate the expression of prepulse inhibition. MolPsychiatry 10, 201–207 (2005).

100. Gill KM, Lodge DJ, Cook JM, Aras S, Grace AA. A novel alpha5GABA(A)R-positive allosteric modulator reverses hyperactivation of the dopamine system in the MAM model of schizophrenia. Neuropsychopharmacology 36, 1903–1911 (2011).

101. Schmid LC, et al. Dysfunction of Somatostatin-Positive Interneurons Associated with Memory Deficits in an Alzheimer’s Disease Model. Neuron 92, 114–125 (2016).

102. Zurek AA, et al. Sustained increase in α5GABAA receptor function impairs memory after anesthesia. J Clin Invest 124, 5437–5441 (2014).

103. Koh MT, Rosenzweig-Lipson S, Gallagher M. Selective GABA(A) alpha5 positive allosteric modulators improve cognitive function in aged rats with memory impairment. Neuropharmacology 64, 145–152 (2013).

104. Engin E, et al. Tonic Inhibitory Control of Dentate Gyrus Granule Cells by alpha5-Containing GABAA Receptors Reduces Memory Interference. J Neurosci 35, 13698–13712 (2015).

105. Taniguchi H, et al. A resource of Cre driver lines for genetic targeting of GABAergic neurons in cerebral cortex. Neuron 71, 995–1013 (2011).

106. Chao HT, et al. Dysfunction in GABA signalling mediates autism-like stereotypies and Rett syndrome phenotypes. Nature 468, 263–269 (2010).

107. Tsien JZ, et al. Subregion- and cell type-restricted gene knockout in mouse brain. Cell 87, 1317–1326 (1996).

108. Cushman JD, et al. Juvenile neurogenesis makes essential contributions to adult brain structure and plays a sex-dependent role in fear memories. Frontiers in behavioral neuroscience 6, 3 (2012).

