## Supplemental material for "Control of contextual memory through interneuronal α5-GABA_A_ receptors"

**Supplementary Information**

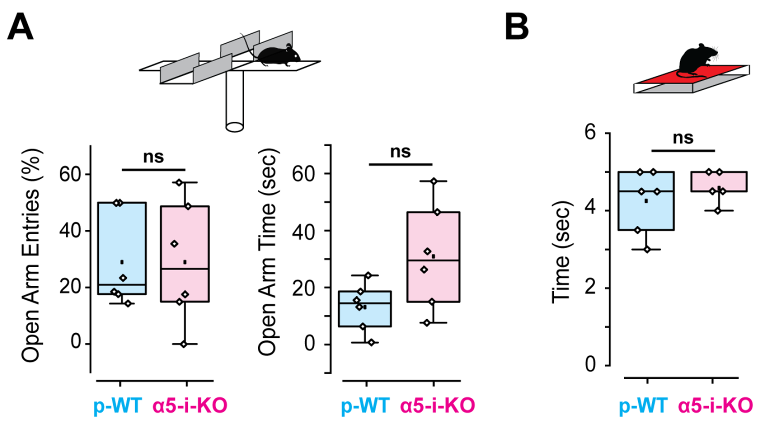


Supplementary Figure 1. Baseline behavioral characteristics of p-WT vs. α5-i-KO mice.

(A) There were no differences in elevated plus maze open arm entries (t(9.31) = 0.000066, p = 0.99, Welch's t test), or open arm entry time (t(6.9) = 2.1, p = 0.073, Welch's t test) between the two genotypes (n = 6 for each), indicating a lack of difference in anxiety.

(B) Latency to paw withdrawal was not different between the two genotypes (n = 6 for each; t(7.0) = 0.91, p = 0.40, Welch's t test), indicating no difference in pain sensitivity. For both parts, each diamond symbol represents one mouse, and data are graphed in quartiles with standard deviation.

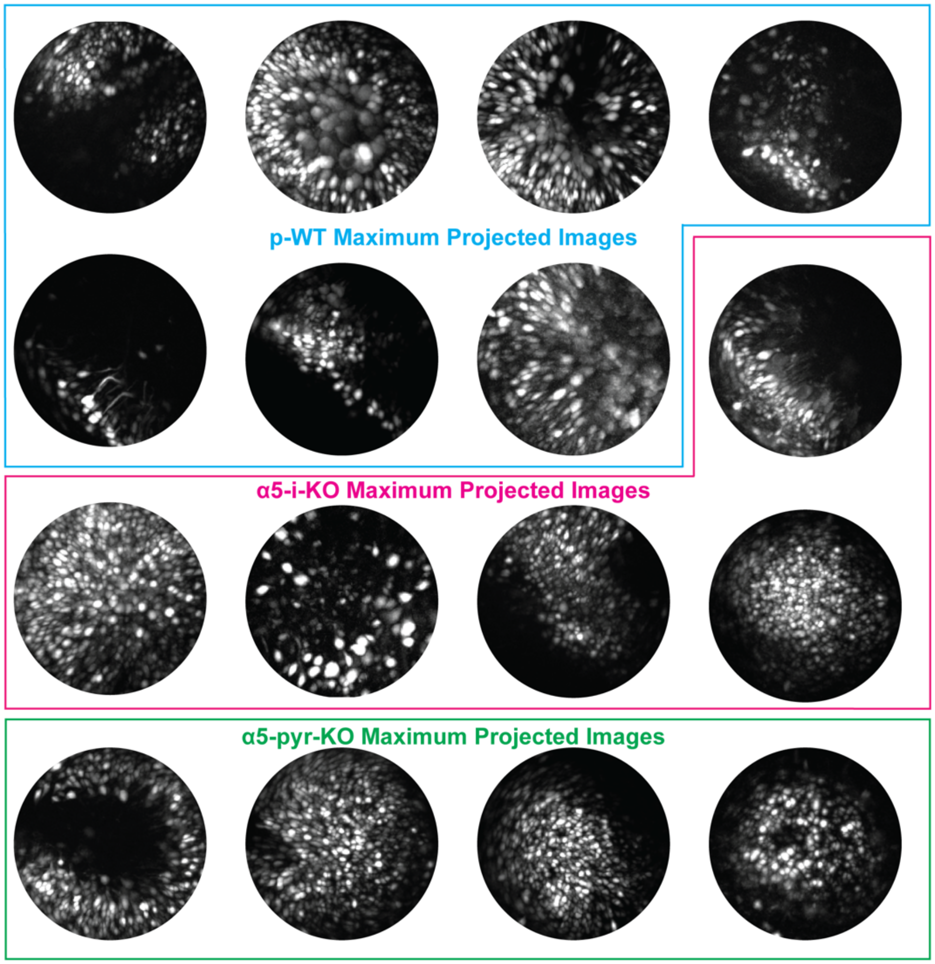


Supplementary Figure 2. Maximum projected images from $Ca^{2+}$ imaging recordings. A range of ~80 to ~800 cells were programmatically captured from CNMF-E algorithm in the first stage of analysis.

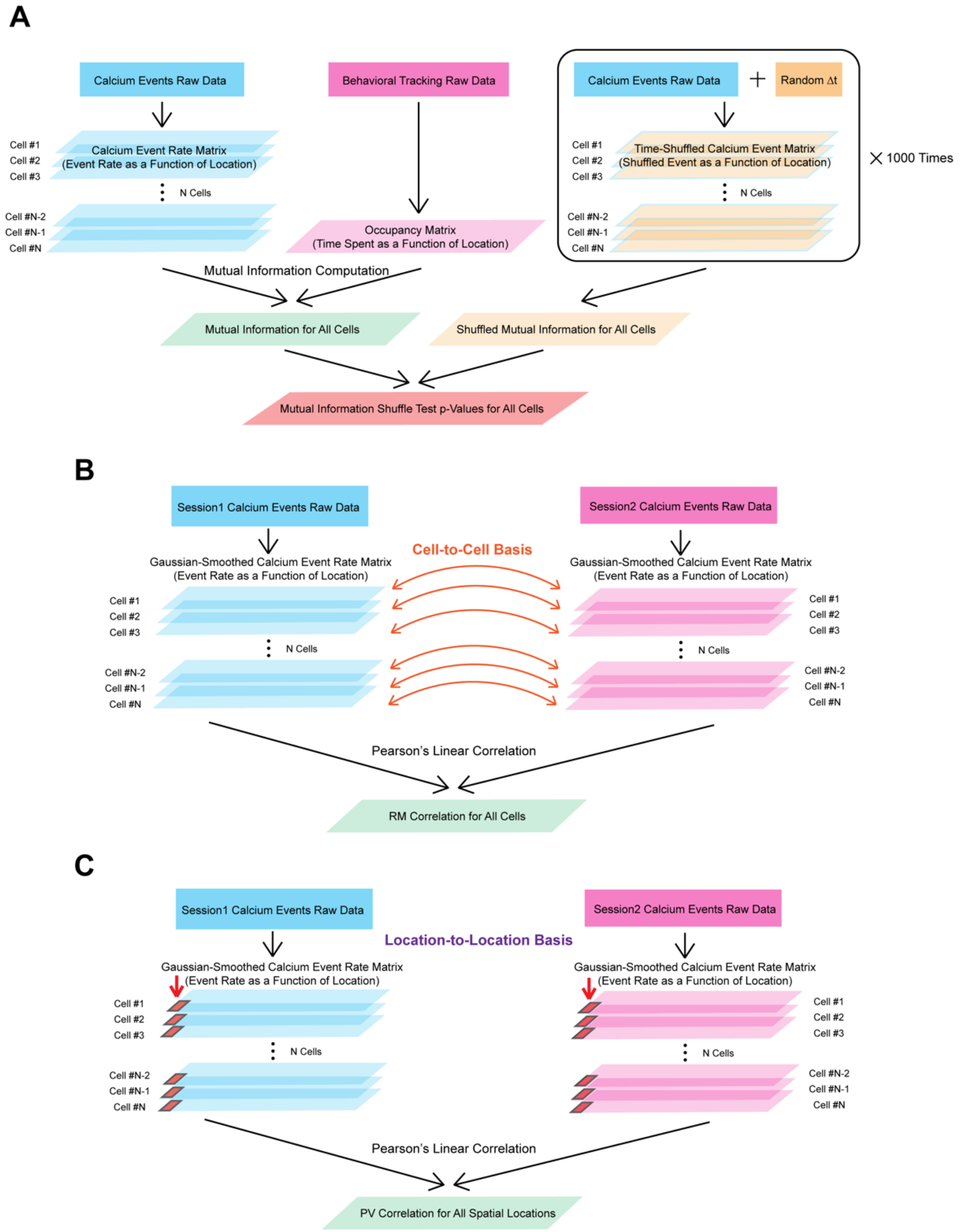


Supplementary Figure 3. Analysis workflow for place cells and spatial engrams.

(A) Calculation of place-specific firing by mutual information (cells with $p(MI)<0.05$ are classified as place cells).

(B) Calculation of cell-based rate map (RM) correlation for both place and non-place cells.

(C) Calculation of position-based population vector (PV) correlation for both place and non-place cells.

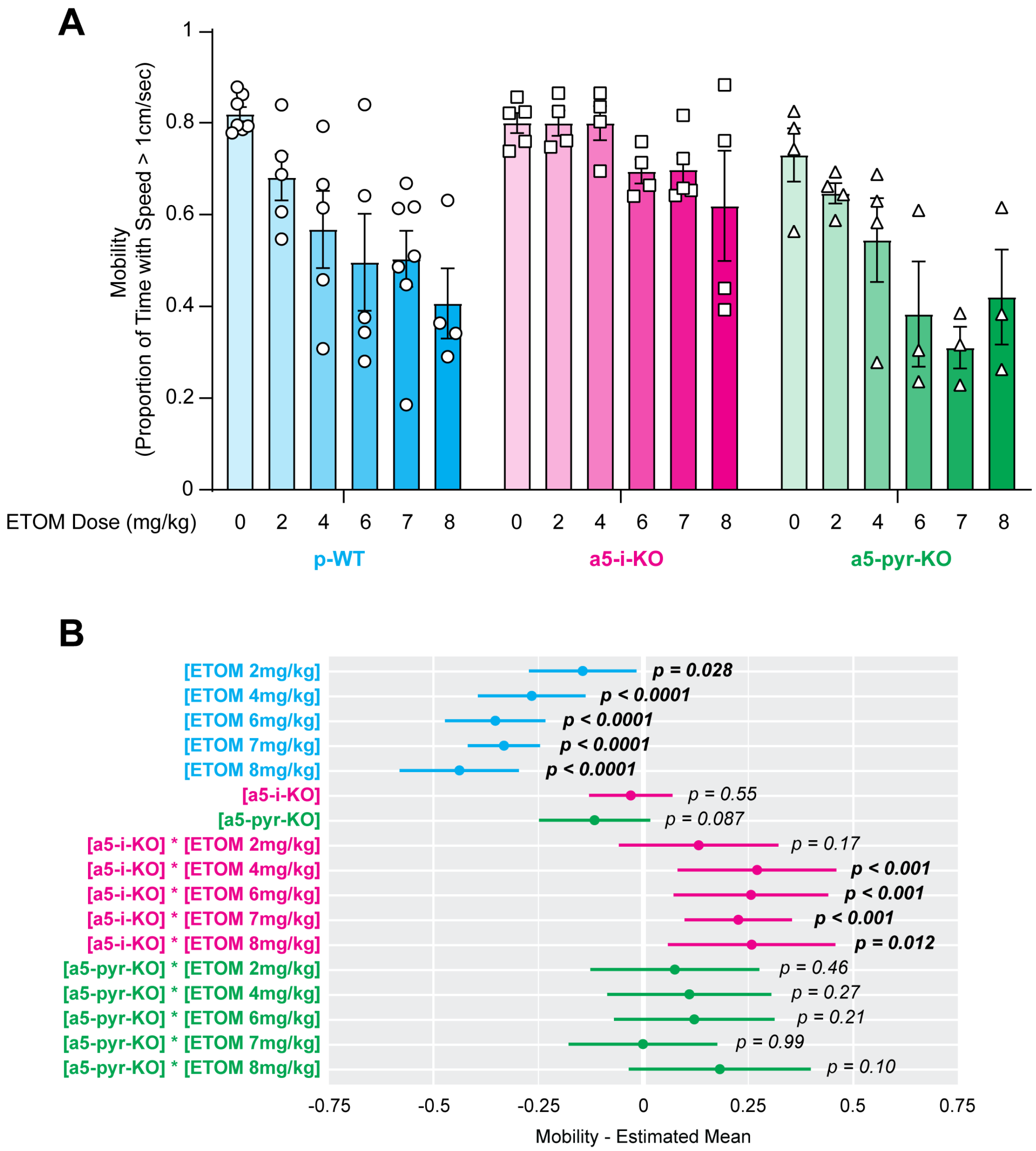


Supplementary Figure 4. Dose-dependent effect of etomidate on mobility.

(A) Fraction of time mice actively explored (± sem) as a function of etomidate dose. Each point represents the mean mobility throughout a 10-min session, for all sessions under a given condition, for each mouse.

(B) A linear mixed effects model was used to evaluate drug effect (first five rows), genotype effect (next two rows), and genotype-drug interactions for mean event rate (next ten rows), using p-WT [genotype] and saline [drug] as reference levels. Etomidate reduced mobility at all doses in p-WT mice. No genotype effect was observed in the reference drug (saline) condition, indicating that elimination of α5-GABA$_A$Rs from interneurons or pyramidal neurons did not alter intrinsic activity. Significant genotype-drug interactions were observed at etomidate doses of 4, 6, 7, and 8 mg/kg for α5-i-KO but not α5-pyr-KO mice, indicating that α5-i-KO mice partially resisted the sedative effect of etomidate.

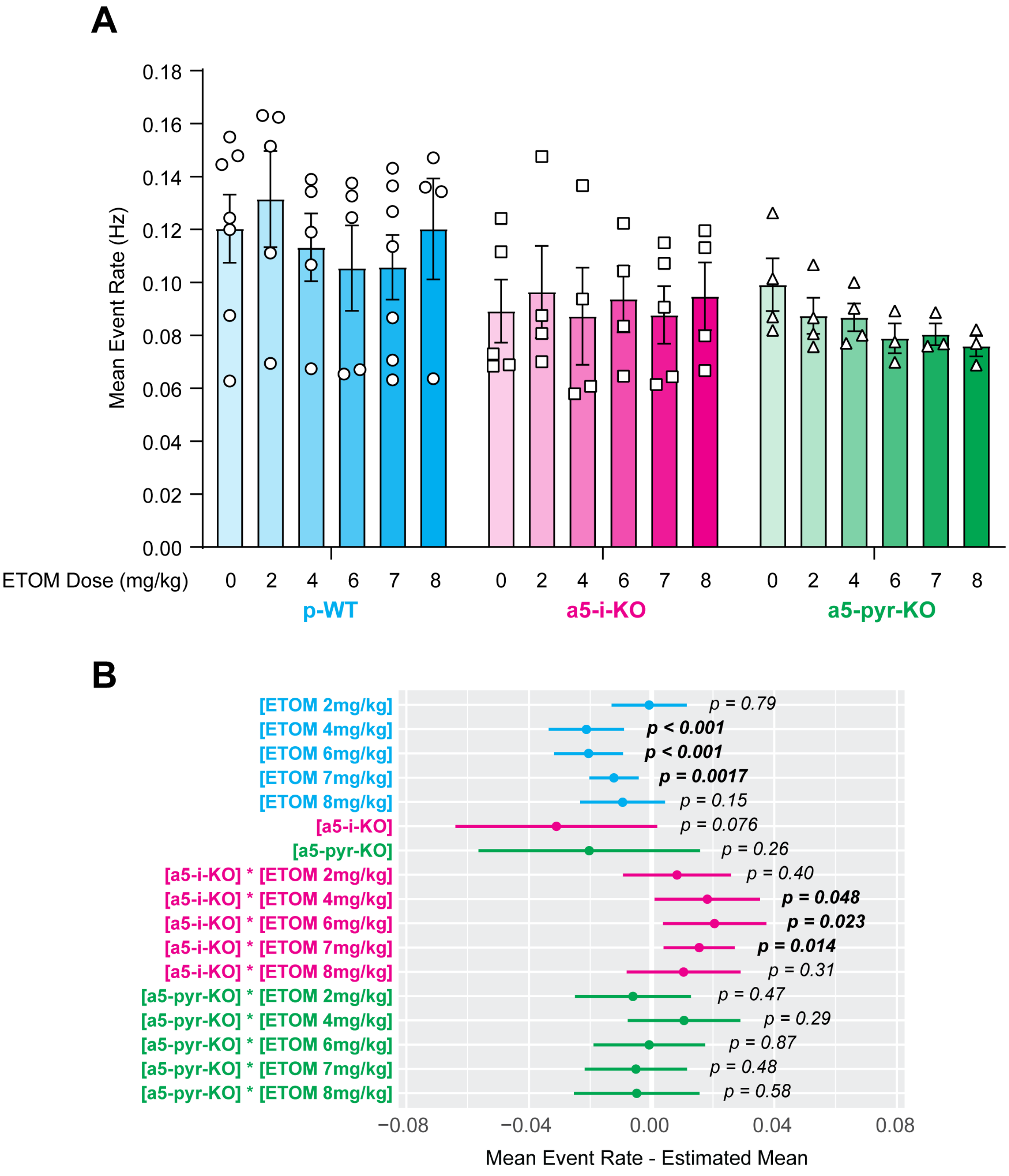


Supplementary Figure 5. Dose-dependent effect of etomidate on mean $Ca^{2+}$ event rate.

(A) Mean event rate (± sem) as a function of etomidate dose for all three genotypes. Each point represents the mean event rate of all cells in all sessions under a given condition for each mouse.

(B) A linear mixed effects model was used to evaluate drug effect (first five rows), genotype effect (next two rows), and genotype-drug interactions for mean event rates (next ten rows), using p-WT [genotype] and saline [drug] as reference levels. Though not visually obvious in part (A), slight (-0.01 to -0.02 Hz) but statistically significant reductions were seen in mean even rate at etomidate doses of 4, 6, and 7mg/kg in p-WT mice. No genotype effect was observed in the reference drug (saline) condition, indicating that elimination of α5-GABA$_A$Rs from either interneurons or pyramidal neurons did not influence cellular activity as reported by $Ca^{2+}$ events. Interestingly, we observed complete resistance to the event rate-suppressing effect of etomidate at doses of 4, 6, and 7mg/kg in α5-i-KO mice, but not in α5-pyr-KO mice.

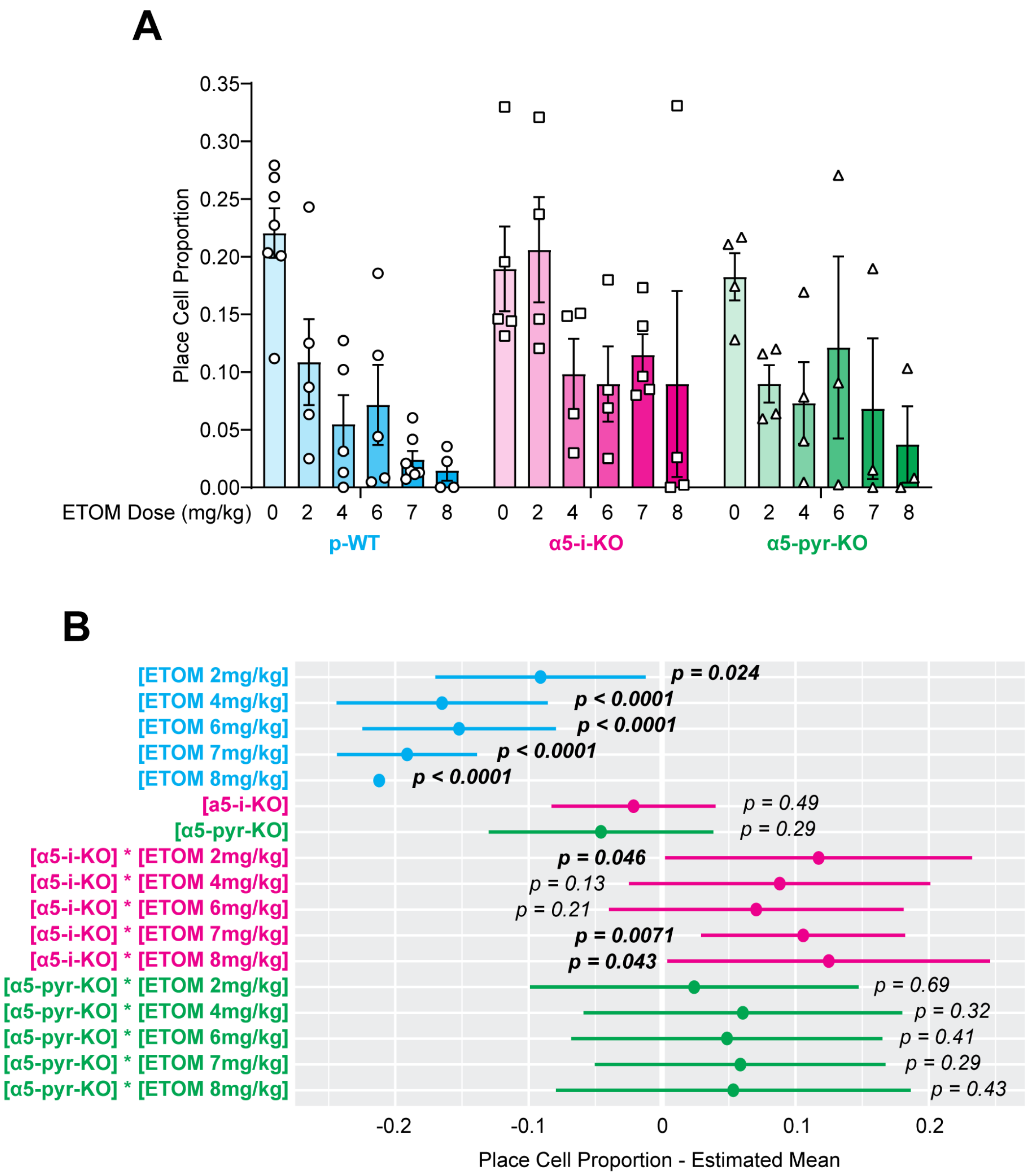


Supplementary Figure 6. Dose-dependent effect of etomidate on place cells.

(A) Mean place cell proportion (± sem) as a function of etomidate dose . Each point represents the mean proportion of all active cells in a session that qualified as bona-fide place cells ($p(MI)<0.05$) for all sessions under a given condition for each mouse.

(B) A linear mixed effects model was used to evaluate drug effect (first five rows), genotype effect (next two rows), and genotype-drug interactions for mean event rate (next ten rows), using p-WT [genotype] and saline [drug] as reference levels. Etomidate strongly suppressed place cell formation at all doses in p-WT mice. No genotype effect was observed in the reference drug (saline) condition, indicating that elimination of α5-$GABA_A$Rs from interneurons or pyramidal neurons did not influence place cell formation under control conditions. Significant genotype-drug interactions were observed at etomidate doses of 2, 7, and 8 mg/kg for α5-i-KO but not α5-pyr-KO mice, implicating interneuronal α5-$GABA_A$Rs as essential targets by which etomidate suppresses spatially modulated firing of pyramidal neurons.

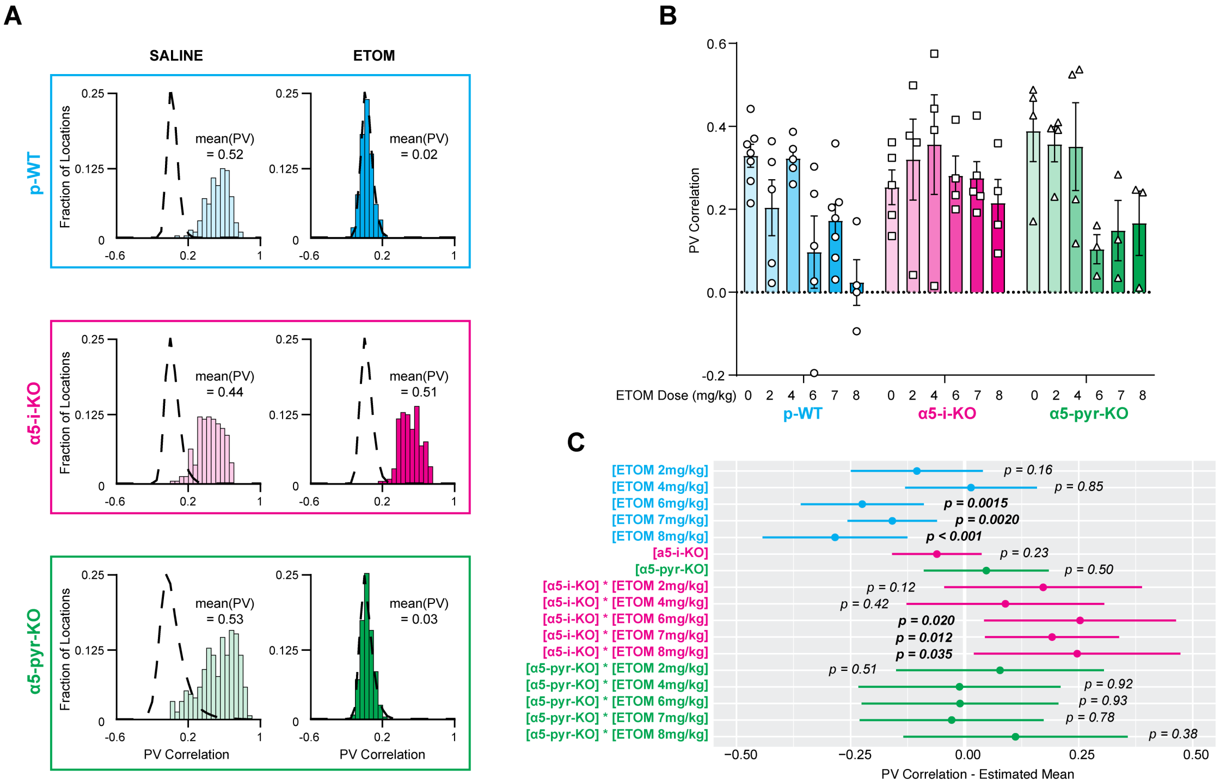


Supplementary Figure 7. PV correlation analysis.

**(A)** Distributions of PV correlations from six paired recording sessions in p-WT (top), α5-i-KO (middle), and α5-pyr-KO (bottom) mice that were administered saline (left) or 7mg/kg etomidate (right). Left: under control conditions, distributions fell substantially to the right of the location-shuffled PV null distributions (dashed lines) in all three genotypes, revealing the presence of stable spatial engrams. Right: etomidate caused the distributions to be shifted toward the null distributions in p-WT and α5-pyr-KO mice but not in α5-i-KO mice.

(B) Mean $PV_{corr}$ (± sem) as a function of etomidate dose. Each point represents the mean $PV_{corr}$ value for all cells active in both of a pair of matched sessions, for all sessions under a given condition for each mouse.

(C) A linear mixed effects model was used to evaluate drug effect (first five rows), genotype effect (next two rows), and genotype-drug interactions for mean event rate (next ten rows), using p-WT [genotype] and saline [drug] as reference levels. Etomidate reduced $PV_{corr}$ at doses of 6, 7, and 8mg/kg in p-WT mice. No genotype effect was observed in the reference drug (saline) condition, indicating that elimination of α5-$GABA_A$Rs from interneurons or pyramidal neurons did not influence $PV_{corr}$ under control conditions. Significant genotype-drug interactions were observed at etomidate doses of 6, 7, and 8 mg/kg for α5-i-KO but not α5-pyr-KO mice, implicating interneuronal α5-$GABA_A$Rs as essential targets by which etomidate suppresses spatial engrams as reported by $PV_{corr}$.

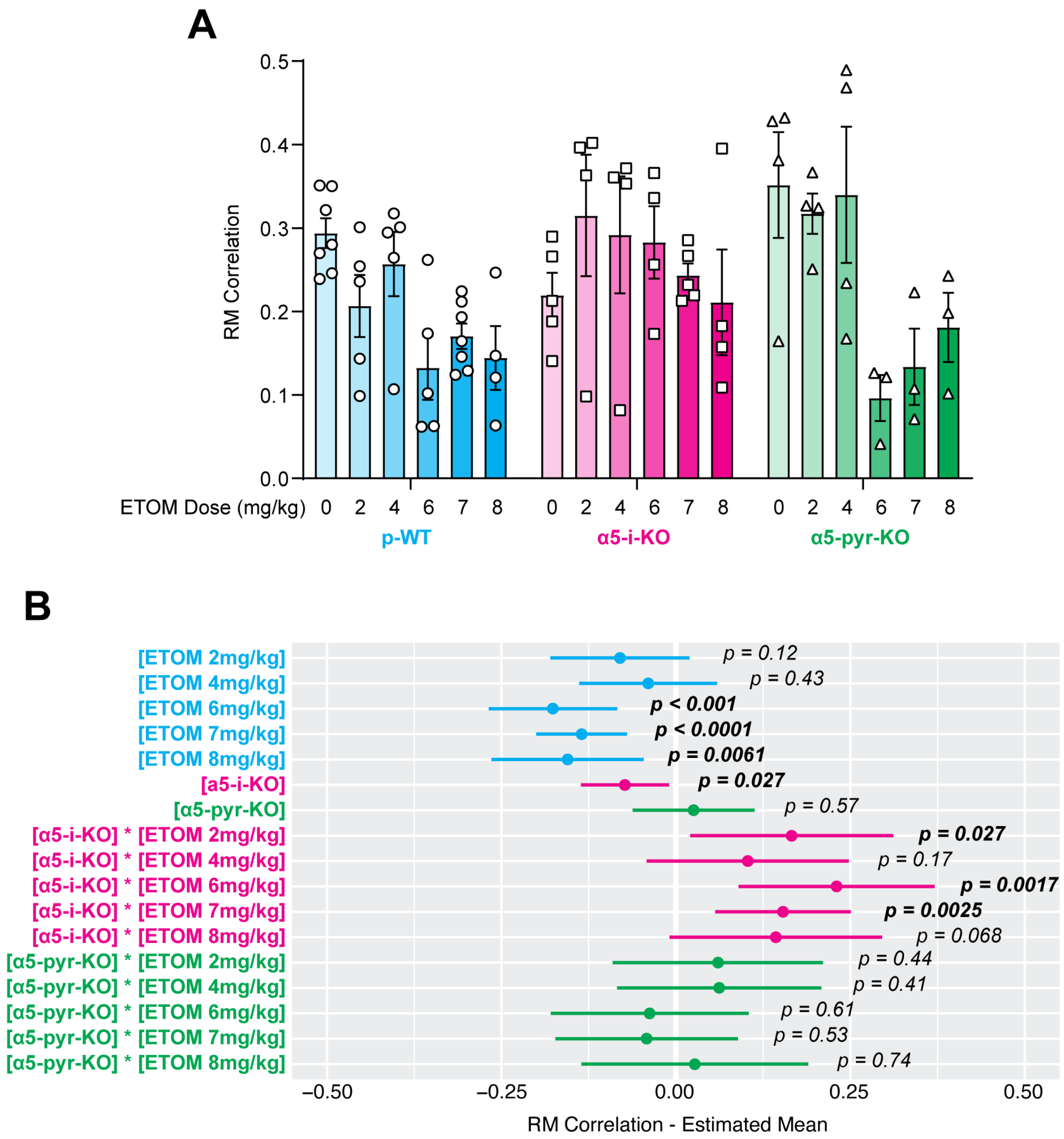


Supplementary Figure 8. Dose-dependent effect of etomidate on RM correlation.

(A) Mean $RM_{corr}$ (± sem) as a function of etomidate dose. Each point represents the mean $RM_{corr}$ value for all cells active in both of a pair of matched sessions, for all sessions under a given condition for each mouse.

(B) A linear mixed effects model was used to evaluate drug effect (first five rows), genotype effect (next two rows), and genotype-drug interactions for mean event rate (next ten rows), using p-WT [genotype] and saline [drug] as reference levels. Etomidate reduced $RM_{corr}$ at doses of 6, 7, and 8mg/kg in p-WT mice. A significant genotype effect in α5-i-KO mice in the reference drug (saline) condition indicates that α5-$GABA_A$Rs in interneurons exert a physiological memory-promoting influence. Significant genotype-drug interactions were observed at etomidate doses of 6, 7, and 8 mg/kg for α5-i-KO but not α5-pyr-KO mice, implicating interneuronal α5-$GABA_A$Rs as essential targets by which etomidate suppresses spatial engrams as reported by $RM_{corr}$.

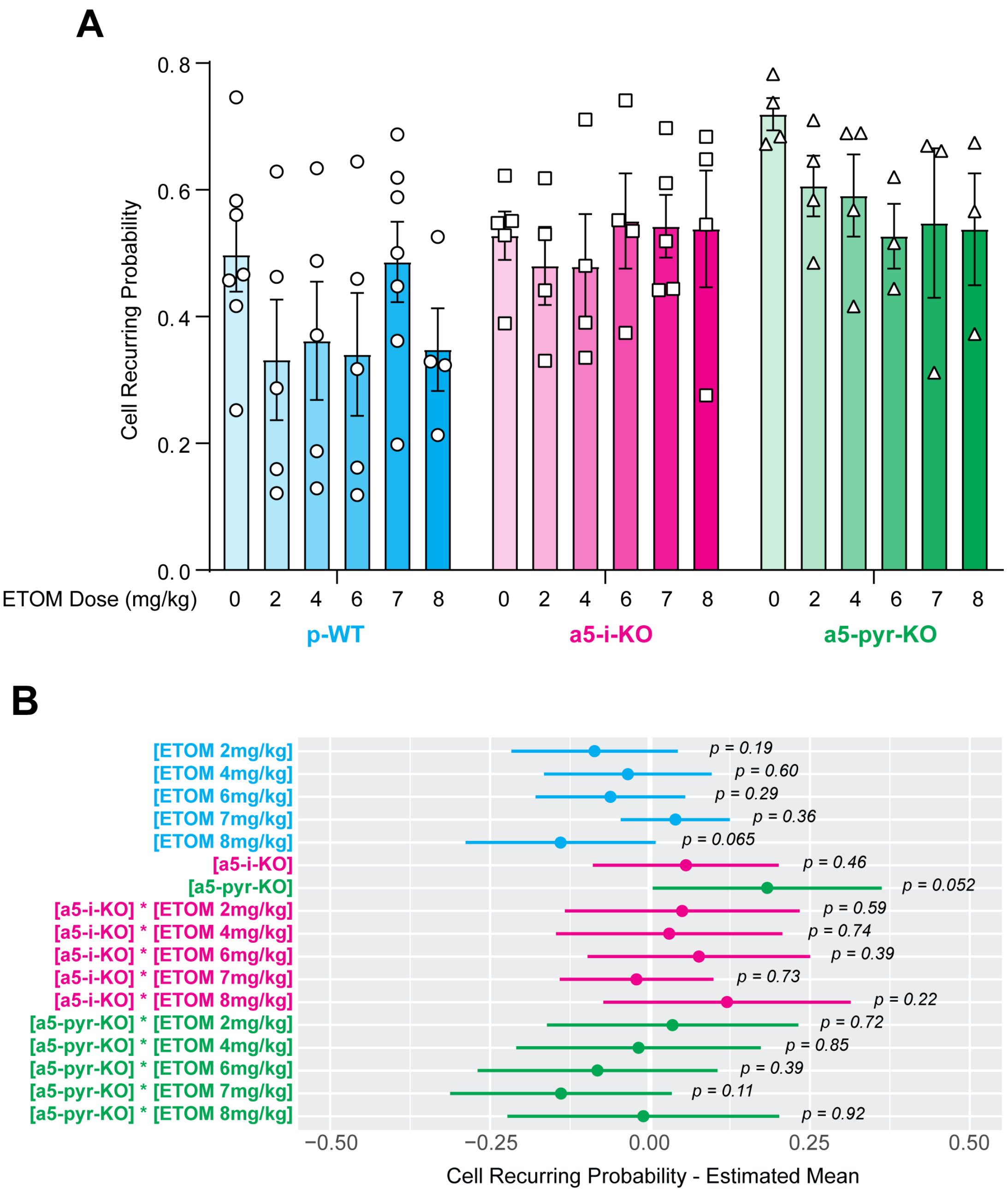


Supplementary Figure 9. Lack of effect of etomidate on cell recurrence probability.

(A) Mean recurring probability (± sem) as a function of etomidate dose. Each point represents the mean recurring probability for all cells active in the AM session for all pairs of matched sessions under a given condition for each mouse.

(B) A linear mixed effects model was used to evaluate drug effect (first five rows), genotype effect (next two rows), and genotype-drug interactions for mean event rate (next ten rows), using p-WT [genotype] and saline [drug] as reference levels. Etomidate did not have any effect on recurring probability at any dose in any genotype.

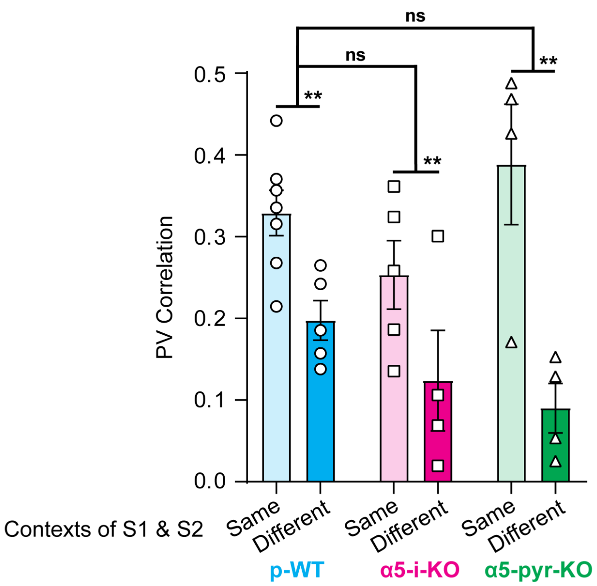


Supplementary Figure 10. Summary of different-context experiments for PV correlation. Similar to RM correlation, in all genotypes, changing contextual cues produced a significant reduction in PV correlation. No genotype-experimental condition interactions were observed, indicating that all three genotypes distinguished contexts equally well.

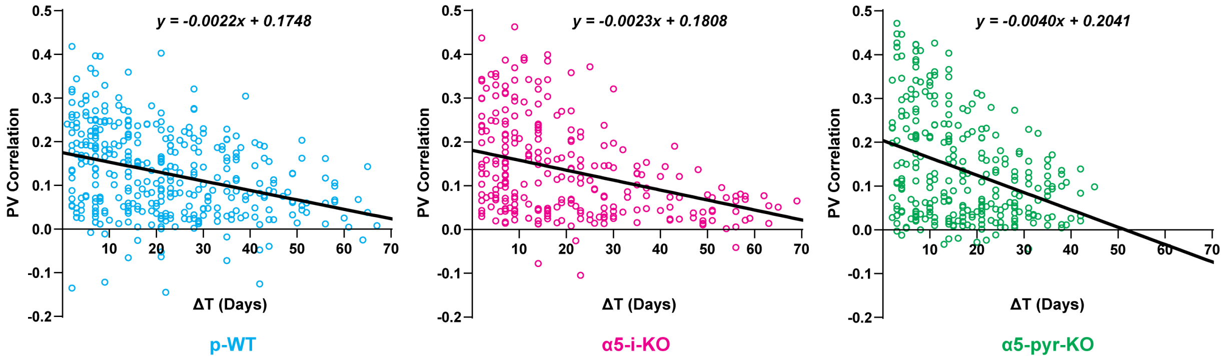


Supplementary Figure 11. Cross-day different contexts experiments analyzed using PV correlation. The conclusions and interpretations were essential identical to that of the RM correlation.

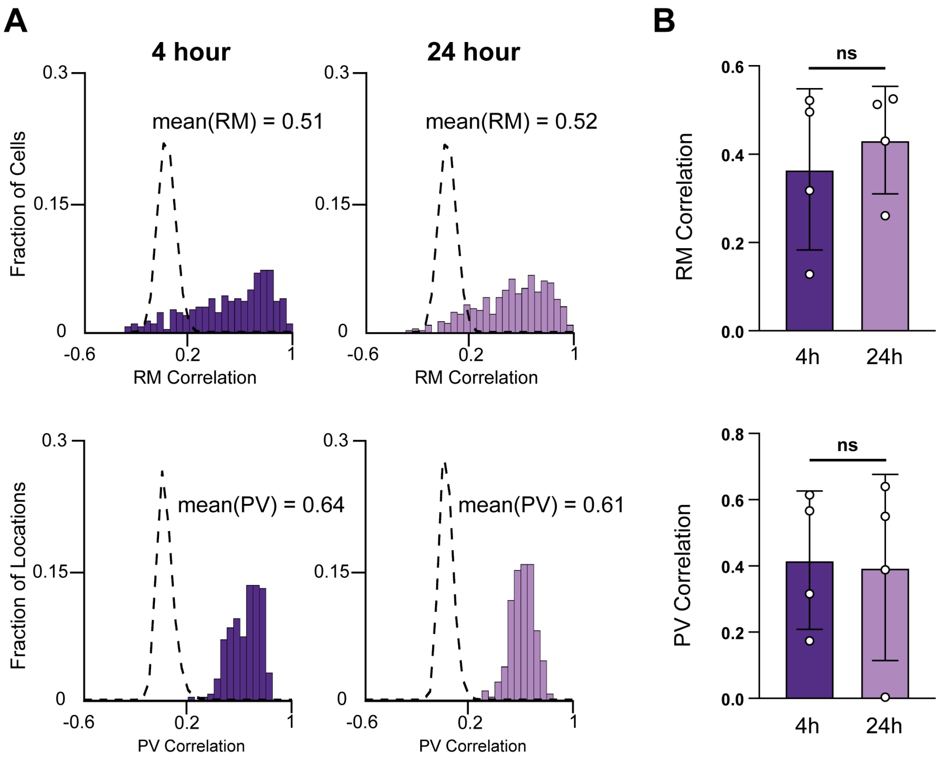


Supplementary Figure 12. 4hr vs 24hr spatial engram stability experiments conducted in a separate cohort of four C57BL/6J mice.

(A) Representative RM and PV correlation distributions when same-context experiments were conducted over 4hrs vs. 24hrs. All four distributions fall significantly outside of their corresponding shuffled null distributions, indicating formation of stable spatial engrams. More importantly, spatial engram stability was essentially identical over 4hrs vs. 24hrs.

(B) Summary of 4hr vs. 24hr experiments, where RM and PV correlations show approximately equal strengths.

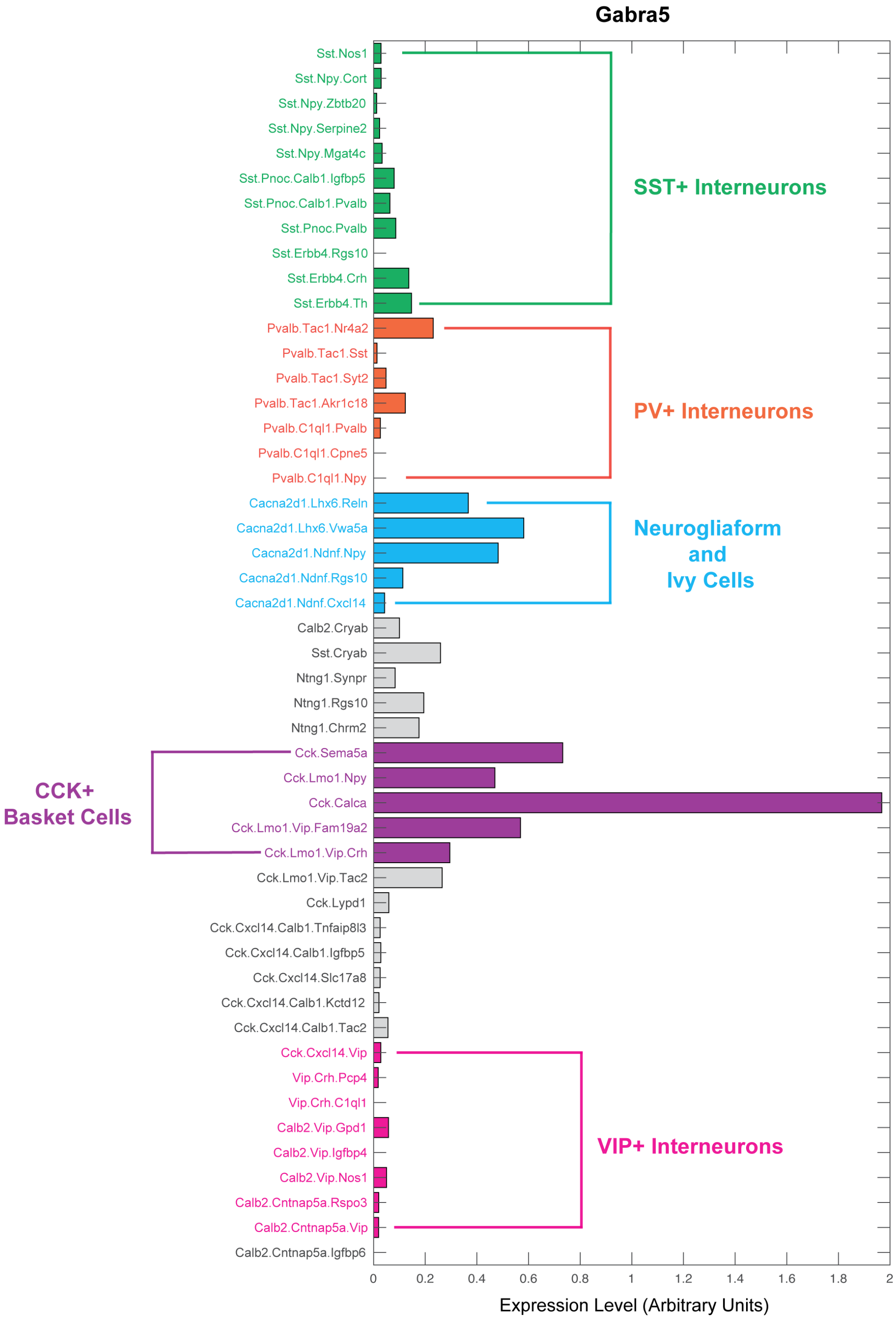


Supplementary Figure 13. Transcriptome profile of Gabra5 mRNA in hippocampal CA1 interneurons. Cell classes are named according to the publication from which the data were obtained [90]. Expression levels were extracted using software provided by the author, together with custom-written MATLAB codes.
